# Immediate and sustained effects of cobalt and zinc-containing pigments on macrophages

**DOI:** 10.1101/2021.04.20.440447

**Authors:** Julie Devcic, Manon Dussol, Véronique Collin-Faure, Julien Pérard, Daphna Fenel, Guy Schoehn, Marie Carrière, Thierry Rabilloud, Bastien Dalzon

**Author notes:** to whom correspondence should be sent or.

## Abstract

Pigments are among the oldest nanoparticulate products known to mankind, and their use in tattoos is also very old. Nowadays, 25% of American people aged 18 to 50 are tattooed, which poses the question of the delayed effects of tattoos. In this article, we investigated three cobalt (Pigment Violet 14 (purple color)) or cobalt alloy pigments (Pigment Blue 28 (blue color), Pigment Green 14 (green color)), and one zinc pigment (Pigment White 4 (white color)) which constitute a wide range of colors found in tattoos. These pigments contain microparticles and a significant proportion of submicroparticles or nanoparticles (in either aggregate or free form). Because of the key role of macrophages in the scavenging of particulate materials, we tested the effects of cobalt- and zinc-based pigments on the J774A.1 macrophage cell line. In order to detect delayed effects, we compared two exposure schemes: acute exposure for 24 hours and an exposure for 24 hours followed by a 3-day post-exposure recovery period. The conjunction of these two schemes allowed for the investigation of the delayed or sustained effects of pigments. All pigments induced functional effects on macrophages, most of which were pigment-dependent. For example, Pigment Green 19, Pigment Blue 28, and Pigment White 4 showed a delayed alteration of the phagocytic capacity of cells. Moreover, all the pigments tested induced a slight but significant increase in tumor necrosis factor secretion. This effect, however, was transitory. Conversely, only Pigment Blue 28 induced both a short and sustained increase in interleukin 6 secretion. Results showed that in response to bacterial stimuli (LPS), the secretion of tumor necrosis factor and interleukin 6 declined after exposure to pigments followed by a recovery period. For chemoattractant cytokines (MCP-1 or MIP-1α), delayed effects were observed with a secretion decreased in presence of Pigment Blue 28 and Pigment violet 14, both with or without LPS stimuli. The pigments also induced persisting changes in some important macrophage membrane markers such as CD11b, an integrin contributing to cell adhesion and immunological tolerance. In conclusion, the pigments induced functional disorders in macrophages, which, in some cases, persist long after exposure, even at non-toxic doses.

**Contribution to the field statement:** Unlike dyes, which are water-soluble, pigments are water-insoluble and thus viewed as inert coloring substances. However, historical pigments such as lead white or vermilion (mercuric sulfide) have been shown to be toxic, suggesting that pigments inertness may not be complete under biological conditions.

Pigments being particulate (nano) materials, they are taken up by professional scavenger cells such as macrophages once they have penetrated into the body. One epitome of this situation is represented by tattooing: the pigments injected with the ink are taken up by dermal macrophages, which life cycle ensures the localized persistence of tattoos over time.

Using an in vitro macrophage culture system adapted to study delayed effects, we have investigated the effects of a series of cobalt- and zinc-containing pigments on macrophages.

First of all, the toxicity of the pigments correlated well with their solubility in acidic media, i.e. conditions prevailing in the phagolysosomes. Even when used at non-toxic doses, the cobalt and zinc pigments showed immediate and/or delayed effects on macrophage functions such as phagocytosis, adhesion, tissue repair or response to bacterial stimuli. Overall, these results show that some pigments may not be as inert as previously thought, and describe a system to investigate these effects.

## 1. Introduction

### 1.1. Definition, prevalence of tattoos and medical complications

Tattooing is an ancient practice which originated in the Stone Age, 6000 years BCE [1, 2]. It consists in the intradermal injection of non-biodegradable and persistent exogenous pigments. Since the 1970, tattoos have been trending and they increasingly appeal to youngsters. In the USA, 25% of people aged 18 to 50 are tattooed. In Europe, the prevalence is approximatively 10% (percentage obtained in Germany and England) [3], meaning that approximately one hundred million people are tattooed in Europe [4]. Other studies indicate that 65% of US tattooed people obtained their first tattoo before age 24. In addition, young people seem to have larger and more tattoos than their elders [5]. However, the insertion of exogenous pigments in the dermis is not without side effects: 67.5% of tattooed people described skin problems comprising short-term side effects (lasting for a few weeks) which are relatively benign and well-documented. They include the classical symptoms of inflammation: discomfort, swelling, and erythema in the tattooed area. However, 6 % of tattooed people report persistent effects linked to tattoos, according to the Klüg’s study [3]. As a comparison, according to Agence Nationale de Sécurité du Médicament et des produits de santé (ANSM, the French drug regulatory agency), an adverse reaction to a product is considered to be frequent when it is found in 1 to 10% of people. It also appears that the effects vary depending on the colors used [4,6–8].

### 1.2. Use and component of tattoo inks

Historically, tattoos are an aesthetic practice corresponding to cultural, ritual or social roles but their applications continue to expand. For example, tattooing is used for cosmetic (permanent makeup) or medical purposes such as scar camouflage, nipple-areola tattooing or colon mapping and oncological radiotherapy (to visualize the location of the area to be treated). These aesthetical and medical applications require different types of inks and colors.

This is why tattoos are currently not limited to black ink (carbon black), and the use of different inks, even though it has increased recently, go back a long way [9]. Thus, tattoos are generally based on a wide range of colored substances in suspension and these mixtures of substances have evolved over time to improve the range of colors or because of national and international regulations [10, 11]. However, mixtures of substances are still not well documented due to the persistent inadequacy of [4, 12] and clandestine tattoos (up to one quarter of all tattoos) [4]. Starting in the 1970s, these pigments were composed of various mineral and organometallic (metal phthalocyanines) in micro and nanoparticulate forms. Gradually, they were officially replaced by organic pigments (e.g. azo pigments) [13] without actually removing metals completely (e.g. cadmium, iron, cobalt, etc.) [4].

### 1.3. Long-term complications of pigments

These factors account for the rise in skin injuries, such as dermatosis or neoplasia, reported in the literature [14, 15]. These short-lived and localized effects (e.g. hypersensitivity skin reactions) are widely known [16]. However, the injection of tattoo inks can also have chronic or long-term effects. Indeed, clinical reports have described cutaneous pathologies at the tattooed zones several years after the tattooing process. These pathologies can be caused by a greater sensitivity to certain bacteria (pathogens or bacteria of the cutaneous flora). Chronic complications linked to sterile inflammation have also been widely described and there are suspicions about systemic effects linked to tattoo inks. The number of case reports describing skin pathology tend to increase with the rise of tattooing practices. They include but are not limited to: allergic reactions, type IV hypersensitivity, autoimmune disorders, spongiotic or atopic dermatitis, lichenoid reaction, local or systemic sarcoidosis. Complications depend on the color and material composing the pigments. For example, red inks can induce a delayed type-IV allergy, black inks can induce chronic inflammations such as granulomatous inflammation or sarcoidosis. [6]. Blue pigments (suspected to contained cobalt) and other colors such as purple or white, could also be linked to sarcoidosis. [4, 7].

Moreover, tattooing seems associated with several cases of neoplasm and skin cancer (e.g. melanoma or carcinoma), which incidence appears to be color- or material-dependent. Red inks (composed of azo pigments), for example, increase UV-induced cancers [8]. Black and dark-blue pigments used in tattoos seem to be linked to the majority of post-tattoo skin cancers. These inks may contain impurities which are carcinogenic substances. Carbon black, for example [17], may contain such impurities as polycyclic aromatic hydrocarbons (PAHs) and phenols [18]. Certain inks contain monoazo pigments (e.g., yellow or red inks) or heavy metals such as mercury sulfide (red), cadmium (from yellow to red), chromium oxide (green), and cobalt (from purple to green), which are also suspected carcinogens [17,19,20]. However, the links between inks and these long-term complications has remain unclear because they were established with descriptions of cases and histological observations. The original studies concerning the understanding of the cellular and molecular mechanisms are still lacking.

### 1.4. Potential role of macrophages in post-tattoo skin pathology?

After intradermal injection, pigments (under micro- or nanoparticulate form) are internalized by three main cells: fibroblasts, neutrophils, and macrophages. Neutrophils are mainly present at the site of injection immediately after the injection due to mechanical injuries comprising blood losses, superficial infections [21], inflammations, etc. caused by the insertion of the needle into the dermis. These short-lived cells (about 24h) are then quickly eliminated by macrophages via the efferocytosis process and are consequently absent from the tattoo site in the long term. Fibroblasts internalize a low quantity of pigments contrary to macrophages which are professional phagocytic cells and ensure the lifelong persistence of tattoo inks in the dermis due to successive cycles of pigment capture, release and recapture. Indeed, dead macrophages are replaced by mostly new monocyte-derived macrophages, or, to a lesser extent, via a local process of self-renewal [22, 23].

Chronic inflammations, autoimmune diseases, and the development of cancers have multiple causes. However, it has been shown that they result from an imbalance between the inflammatory and tissue-repair responses implemented by innate immunity, in particular by pro-inflammatory M1 macrophages and alternative (or rather, anti-inflammatory) M2 macrophages. A disruption of the proportion of M1 and M2 macrophages and/or their functionality also has an impact on the adaptive immune response. [24–26]. As an example, studies described a higher density of M2 macrophages in sarcoidosis and atopic dermatitis lesions in comparison to healthy patients [27–30]. Regarding atopic dermatitis, it has been observed that phagocytes’ chemotactic and phagocytosis capacities are deficient [31], leading to an increase in skin infections [30].

### 1.5. Rationale for the study design – importance of testing macrophages functionalities

In this article, we decided to assess the persistent effects of some pigments used in tattoo inks on macrophages, in comparison to the immediate effects, because of their key role in the persistence of tattoos [22, 23]. Because of the versatile roles of macrophages in tissue (skin) homeostasis (the elimination of bacteria, efferocytosis, and the regulation of many other cells via the release of cytokines and chemokines), an impairment of their functionalities could damage tissues and cause other diseases [32, 33]. Furthermore, several articles showed that tattoo ink pigments can be found in lymph nodes due to both passive transport and active transport via macrophages [34]. This is especially true several weeks after the injection of tattoo ink when around 30% of the ink has been eliminated via blood or lymphatic circulation [35–37]. However, the presence of pigment-loaded macrophages in lymph nodes can be due to the migration of particle-loaded macrophages from the tattoo site to the draining lymph nodes, as suggested by some authors in the case of other mineral particles [38, 39]. Whatever the mechanism, a massive absorption of pigments by macrophages might change their functions and thus, alter immune homeostasis at the tattoo site and at distal sites such as lymph nodes.

The present article investigates cobalt and zinc pigments which constitute a wide range of colors found in tattoos (from purple to green and white) [40, 41]. Although the exact nature of the pigments used in tattoo inks is not disclosed, as it is considered to be a trade secret, we chose to test three cobalt-based pigments referenced in the Colour Index International as PG19 (dark green), PB28 (blue), and PV14 (purple), and one zinc-based pigment referenced as PW4 (white). However, cobalt is both an essential element (given that it is a cofactor of vitamin B12) and a toxic one at high doses [42]. It has deleterious effects on fibroblasts [43], leukocytes [44], and macrophages [45, 46], as well as a long-term carcinogenic potential linked to a slow release [47]. Thus, a progressive release of cobalt from pigments internalized by the macrophages may lead to an alteration of their functions and phenotypes.

In low quantities, zinc is an essential trace element and the recommended dietary allowance is between 8 and 11 mg/day for adults. Moreover, zinc plays an important role in numerous biological functions (it is a co-factor in enzymatic reactions, the metabolism of vitamin A, and immunoregulation) [48]. However, an excessive absorption of zinc is considered to be toxic [49]. Zinc oxide nanoparticles are already known to have a toxic effect on immune cells such as macrophages [50]. Toxicity can be explained by the dissolution of these nanoparticles in cells which release zinc ions [51, 52]. This mechanism may be the cause of the potential long-term adverse effects of cobalt and zinc-based pigments, a fact which justifies the relevance of the present work. To this end, we used an in vitro murine macrophage system in load and recovery mode in which the cells are first loaded with particles over 24 hours and then cultured without adding particles for three days [53, 54].

## 2. Materials and methods

### 2.1. Pigment characterization

#### 2.1.1. Pigment description

Pigment Blue 28 (cobalt aluminate, CoAl_2_O_4_), Pigment Violet 14 (cobalt phosphate) and Pigment White 4 (zinc oxide) were purchased from Kama Pigments (Montreal, Canada). Pigment green 19 (mixed cobalt zinc oxide, deep cobalt green CO804) was purchased from Tokyo Pigment (Tokyo, Japan). Corundum (aluminum oxide spinel) was purchased from Merk (Saint-Quentin-Fallavier, France).

Pigments were received as powders. They were suspended in gum arabic (100 mg/ml) that had been previously sterilized overnight at 80°C, with gum arabic thus being used as an anti-agglomeration reagent. Then, the suspension of pigments (100 mg/mL) was sterilized overnight again before being was sonicated in a Vibra Cell VC 750 sonicator equipped with a Cup Horn probe (VWR, Fontenay-sous-Bois, France) with the following settings: time = 30min – 1 second ON, 1 second OFF – Amplitude = 60%, corresponding to 90 W/pulse.

#### 2.1.2. Dynamic Light Scattering (DLS)

The pigments were diluted to 100 µg/ml with water or culture medium and they transferred in UVette^®^ 220-1600 nm (catalog number: 0030106.300, Eppendorf, Montesson, France). Then, the size and dispersity of pigments were measured via Dynamics Light Scattering (DLS) using Wyatt Dynapro Nanostar (Wyatt Technology, Santa Barbara, CA, USA).

#### 2.1.3. Transmission Electronic Microscopy (TEM)

Negative Stain On Grid Technique (SOG): 10 µL were added to a glow discharge grid coated with a carbon supporting film for 3 minutes. The excess solution was soaked off using a filter paper and the grid was air-dried. The images were taken under low dose conditions (<10 e-/Å2) with defocus values comprised between 1.2 and 2.5 μm on a Tecnai 12 LaB6 electron microscope at 120 kV accelerating voltage using CCD Camera Gatan Orius 1000. The measurement was processed with Image J.

### 2.2. Cell culture and viability assay

The mouse macrophage cell line J774A1 was purchased from the European Cell Culture Collection (Salisbury, UK). For cell maintenance, cells were cultured in DMEM containing 10% of fetal bovine serum (FBS) and 5 µg/ml of Ciprofloxacin. Cells were seeded every 2 days at 250,000 cells/ml and harvested at 1 million cells/ml.

For treatment with pigments, the following schemes of exposure were used (Figure 1):

1. An acute (24 hours) exposure with pigments: cells were seeded in 12-well adherent or non-adherent plates (according to the test) at 1 million cells/ml in DMEM containing horse serum 1% (HS 1%) and 5 µg/ml of Ciprofloxacin. Cells were exposed to pigments on the following day and analyzed at the end of the 24h-period of exposure to pigments.
2. A recovery exposure for 4 days: Cells were seeded in 12-well adherent plates at 1million cells/ml in DMEM supplemented with horse serum, as described above. After 24 hours of exposure to pigments, the medium was changed in order to remove pigments. The fresh medium was left for another 48 hours and changed again for 24 hours before cell harvest The choice of a medium supplemented with a low serum concentration was made in order to limit cell proliferation during the recovery period [55]. Indeed, cell proliferation during recovery has been observed before [53] and is not likely to reflect the in vivo situation.

**Figure 1.**
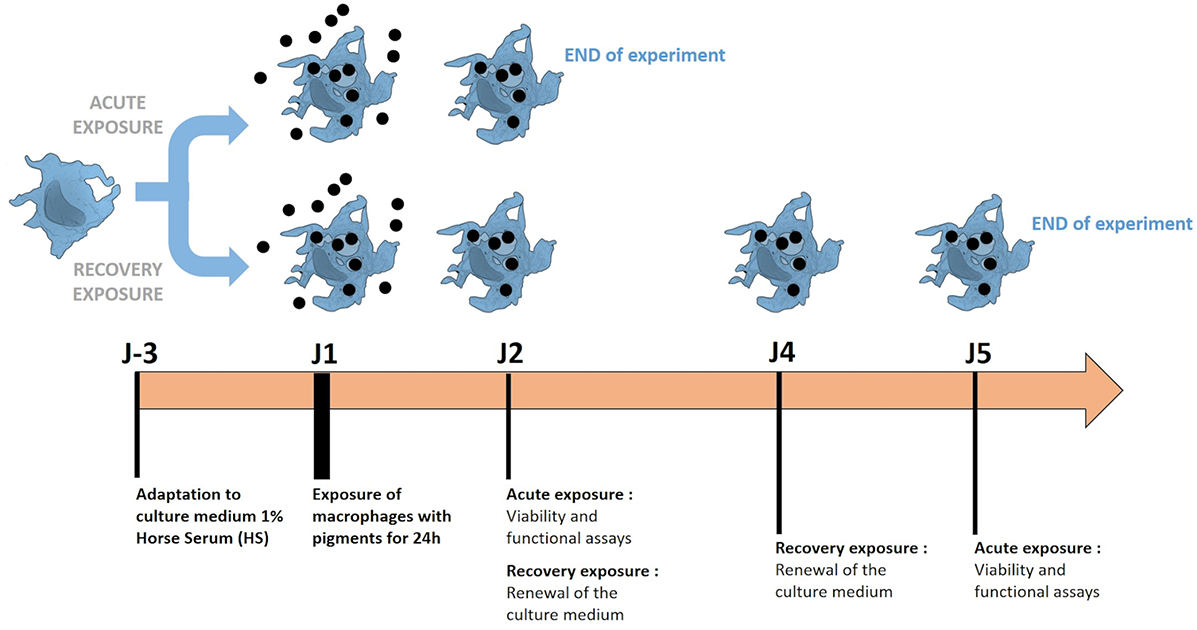
Schematic representation of experiments using macrophages exposed with pigments. The methodology of acute exposure and recovery was illustrated. Cells are grown in a low serum culture medium (1% horse serum), to which they are first adapted. The exposure and recovery phases are then performed to analyze the induced effects.

**Figure 2.**
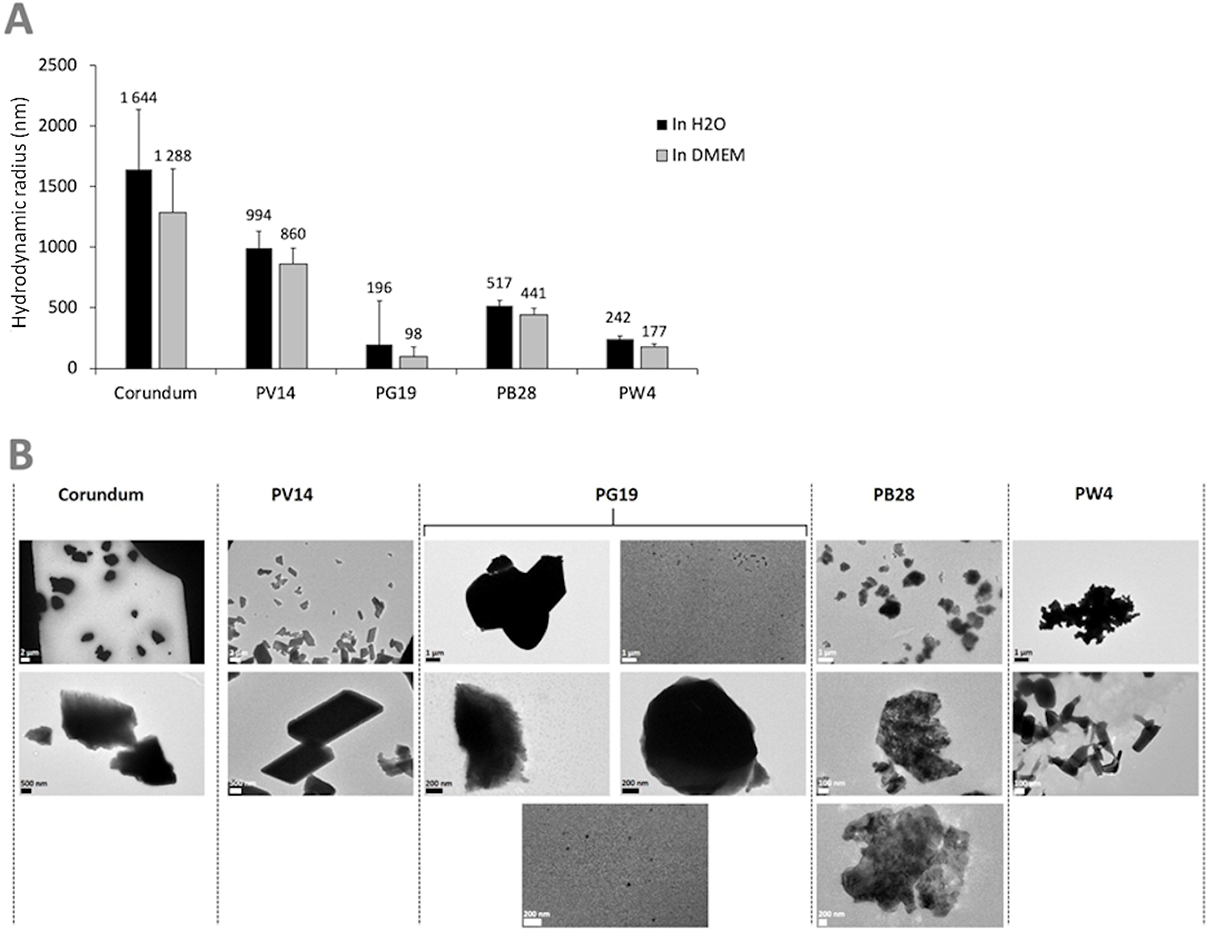
Characterization of pigments PV14, PG19, PB28, PW4 and their corundum control. Panel A: Hydrodynamic diameter measurement by DLS in H_2_O (in black) and DMEM culture medium (in grey). Panel B: visualization of pigments by TEM microscopy in H_2_O.

After exposing the cells to various concentrations of pigments, the viability of the cells was tested. Cells were flushed and harvested with PBS and centrifuged at 200 g for 5min. Pellets were resuspended in PBS with propidium iodide (PI) at a final concentration of 1 µg/ml. The percentage of viability was measured via a FacsCalibur flow cytometer equipped with the CellQuest software program 6.0 (Becton Dickinson Biosciences, Le Pont-de-Claix, France) and settings corresponding to PI (480 nm excitation and 600 nm emission). For every pigment tested, the LD20 was determined.

### 2.3. Measurement of pigment Dissolution via inductively coupled plasma atomic emission spectroscopy

#### 2.3.1. In vitro dissolution (in acidic medium)

Pigments at just below the Lethal Dose 20 (LD 20) concentration for J774A1 macrophages, i.e. 2.5 µg/ml and 500 µg/ml, were mineralized in aqua regia (Mixture of nitric acid and hydrochloric acid, optimally in a molar ratio of 1:3). Samples were diluted in 10% nitric acid prior to analysis by inductively coupled plasma atomic emission spectroscopy (ICP-AES) on a Shimadzu ICP 9000 with Mini plasma Torch instrument used in axial reading mode. A standard range of cobalt, zinc and aluminum (from 3.9 µg/L to 10 mg/L) was prepared for quantification.

#### 2.3.2. In vivo dissolution (in macrophages)

Cells were seeded in 6-well plates at 1 million cells/ml in DMEM containing 1% horse serum and 5 µg/ml Ciprofloxacin. They were exposed to nanoparticles on the following day. For recovery exposure, the medium was changed to remove the pigments and left for another 48 hours. The medium was changed one more time and the cells were left for 24 hours in culture medium. After 24 h (acute exposure) and 96 h (recovery exposure). Following medium removal, the cells were rinsed with PBS 1X and then lysed with 600 µl lysis buffer (HEPES 5 mM, spermine tetrahydrochloride 0.75 mM, tetradecyldimethylammonio propane sulfonate 0.1%). 50 µl of the crude lysate were collected and total amount of protein was quantified with a modified dye-binding assay [56]. The remaining lysate was centrifuged at 10,000 g for 10 minutes to sediment the nuclei and the undissolved pigments. Finally, the supernatant was collected. Samples were kept at −20°C until mineralization.

The samples were thawed and mineralized by the addition of one volume of Suprapure 65% HNO_3_ and incubated on a rotating wheel at room temperature for 18 h. The samples were then diluted up to 6 ml with 10% nitric acid and the metal concentrations were measured as described above. The results were normalized by measuring the protein concentration (used as an indicator for cell number) in the samples.

### 2.4. Assessment of the internalization capacity of macrophages

#### 2.4.1. Measurement of the phagocytic activity

##### a) Quantitative measurement via flow cytometry measurement

The phagocytic activity was measured as previously described [57, 58] using fluorescent latex beads (solid = 2.5%, 1 µm diameter fluorescent yellow-green, catalog number L4655 from Merk). The green fluorescence internalized by the cells allows to quantify the phagocytic activity. Cell culture was performed as described above. Cells were seeded in 12-well plates and exposed to pigments (acute or recovery exposure) at 37°C, 5% CO_2_. The fluorescent beads were coated in a mixture of horse serum 50% and PBS 1X 50% for 30min at 37°C (2.2 µL of fluorescent latex beads for 1 ml). 100 µL/ml of this solution were added to cell culture (final concentration of beads = 5µg/mL), and then, cells were incubated for 3 hours at 37°C. Cells were harvested, washed with PBS 1X, and centrifuged for 5 min at 1200 rpm. The pellets were suspended with 400 µl in PBS1X – propidium iodide 1 µg/ml and analyzed by flow cytometry.

##### b) Qualitative visualization via confocal microscopy

Cells were seeded at 100,000 cells/mL in DMEM+1% horse serum medium on a 22×22mm glass slide previously sterilized. After acute or recovery exposure of cells with pigments, The yellow-green fluorescent beads L4655 (previously coated in a mixture of horse serum 50% and PBS 1X 50% for 30min at 37°C) are added at a final concentration of 5 µg/mL. Then, cells were incubated for 3 hours at 37°C. The actin staining step in order to visualize the cell contour and the nucleus staining step was performed based on [59]. Cells were washed twice for 5 min in PBS, fixed in 4% paraformaldehyde for 30 min at room temperature. Then, after two washes, cells were permeabilized in 0.1% Triton X100 for 5 min at room temperature. Two washed are again performed and Phalloidin-Atto 550 (catalog number : 19083, from Merk) at 250 nM final concentration was added to the cells and let for 20 min at room temperature in the dark. Coverslips-attached cells were washed again and colored 5 min with Dapi 1µg/mL final concentration. After two rinses with PBS 1X, the glass slides were placed on microscope slides (Thermo Scientific, Waltham, MS, USA) using a Vectashield mounting medium (Catalog number : H-1000, from Eurobio, Paris, France) and imaged using a Zeiss LSM 800 confocal microscope (Zeiss, Jena, Germany). The images were processed using the ImageJ software.

#### 2.4.2. Measurement of the lysosomal integrity via Neutral red assay

This stain dyes lysosomes via active transport linked to the proton gradient, and is thus a good indirect assay for lysosomal integrity and function. Cells were seeded in adherent 12-well plates and exposed to pigments (acute or recovery exposure) at 37°C, 5% CO_2_. 2 mg/ml neutral red in EtOH 50% were prepared and 10 µl of this solution were added to every well containing 1 ml of culture medium and incubated for 2 hours at 37°C. After incubation, cells were washed twice with PBS and eluted with 2 ml of acid-alcohol solution (acetic acid 1% v/v, ethanol 50% v/v). The plates were incubated for 30min at room temperature on a rotating table. The red color released was read at 540 nm [60]. To normalize the results, the number of adherent cells was estimated via a crystal violet assay. The fixed cell layer was rinsed twice with PBS and stained with crystal violet (4 µg/ml in PBS) for 30 minutes. Then, the cell layer was rinsed twice with PBS before elution with 1 ml of acid-alcohol solution. The purple color released was read at 590 nm using a spectrophotometer. The 540 nm/590 nm absorbance ratio was used as an index for the lysosomal functionality of the cells.

#### 2.4.3. Visualization of the internalization of pigments in macrophages at a low concentration of exposure (2.5 µg/mL)

Cells were seeded at 100,000 cells/mL in DMEM+1% horse serum medium on a 22×22mm glass slide previously sterilized. After acute or recovery exposure of cells with pigments made fluorescent via linkage with Rhodamine isothiocyanate (RITC). The protocol of mineral particle labelling is described by Dechézelles J-F *et. al.* [61]. For the actin staining step in order to visualize the cell contour, the protocol looks like the one described in part 2.4.1 – b) “microscopic visualization” with some modifications. The actin staining was performed via phalloidin-Atto 390 (catalog number: 50556, Merk) at 1 µM final concentration for 45 min at room temperature in the Dark. The nuclear cell was stained with Sytox red dead cell stain at 50 nM final concentration (catalog number: S34859, ThermoFisher Scientific).

### 2.5. Assessment of the inflammatory capacity of macrophages

#### 2.5.1. Measurement of Nitric oxide (NO) production

The cells were grown to confluence in 12-well plates (harvested at 1 million cells/ml). Half of the wells were treated with 200 ng/ml LPS, and arginine monohydrochloride was added to all the wells (5 nM final concentration) to give an unlimiting concentration of substrate for NO synthase. After 18 hours of incubation, the cell culture medium was recovered, centrifuged at 15,000 g for 15min to remove pigments, non-adherent cells and debris. 500 µL of supernatant were collected. After the addition of an equal volume of Griess reagent, the mixture was incubated at room temperature for 20min in order to wait for the completion of the reaction. The absorbance was read at 540 nm. To normalize the results, the number of cells in the culture plate was estimated via a crystal violet assay. The cell layer was rinsed twice with PBS and fixed with an acid-alcohol solution (acetic acid 1 % v/v, ethanol 50 %v/v) for 30 minutes. The cell layers were then stained with crystal violet as described in section 2.4.2.

#### 2.5.2. Measurement of cytokines and chemokines secretion

Tumor necrosis factor (TNFα) and interleukin 6 (IL-6), macrophage Inflammation Protein (MIP-1α) and monocyte chemoattractant Protein-1 (MCP-1) were measured in the supernatant from cultured cells exposed to pigments and/or activated by LPS 200 ng/ml, using the Mouse Inflammation Cytometric Bead Array kit (reference: BD Biosciences, Rungis, France) according to the manufacturer’s instructions. Measurements were performed on a Facscalibur flow cytometer, and the concentrations of cytokines secreted were assessed using FCAP Array Software.

### 2.6. Measurement of intracellular glutathione

The level of intracellular glutathione were assessed using the monochlorobimane methodology, with some modifications [62]. Cells were harvested, centrifuged during 5 min at 1200 RPM, and labeled with 75 μM monochlorobimane (previously diluted in warm PBS) for 5 min at 37°C. The staining was stopped via an incubation at 4°C (in ice) for 5 min in the dark. Finally, cells were washed twice with cold PBS and the fluorescence were measured via BD FACSMelodyTM flow cytometer (BD Biosciences, Le Pont-de-Claix, France) using a laser excitation at 405 nm and an emission at 448 ± 45 nm.

### 2.7. Measurement of reactive oxygen species

After acute or recovery exposure, culture medium was removed and replaced with warm PBS-DHR 123 (final concentration = 500 ng/mL) (Sigma-Aldrich Catalog number D1054). Then cells were incubated 20 min at 37°C, 5% CO2. After the incubation, cells were harvested in FACS tube previously filled with cold PBS-Glucose (1g/L). Tubes were centrifuged 5 min at 1200 RPM. After the washed step, the supernatants were removed and cell pellets were ressuspended in 400 µL of cold PBS-Glucose (1g/L) containing Sytox red 1/1000 (Catalog number: S34859, ThermoFisher scientific). The fluorescence of DHR123 were measured via Facscalibur flow cytometer (BD Biosciences, Le Pont-de-Claix, France) using a laser excitation at 488 nm and an emission at 530 ± 30 nm. The dead cells were excluded of the analysis using a laser excitation at 630 nm and an emission at 633 ± 670 (long pass filters).

### 2.8. DNA Damage measurement via Comet assay

The presence of strand breaks in the DNA of the cells exposed to the particles was evaluated using the alkaline version of the comet assay, as described [63]. After exposure to particles, cells were collected using trypsin and stored at −80°C in freezing buffer (85.5 g/L sucrose, 50 ml/L DMSO prepared in citrate buffer (11.8 g/L), pH 7.6). Ten thousand cells were mixed with 0.6% low melting point agarose (LMPA) and tw0o drops of this mixture were put on a slide that had previously been coated with agarose. The cell/LMPA mixture was left to solidify on ice for 10 min, and then slides were immersed in cold lysis buffer (2.5 M NaCl, 100 mM EDTA, 10 mM Tris, 10% DMSO, 1% Triton X-100, pH10) and incubated overnight at 4°C. Slides were then rinsed with PBS, and DNA was left to unwind for 30min in the electrophoresis buffer (300 mM NaOH, 1 mM EDTA, pH > 13). Electrophoresis was performed in a vertical electrophoresis tank (Cleaver Scientific) at 0.7 V/cm for 30min. Slides were then neutralized in PBS and stained with 50 µL of GelRed (Thermofisher Scientific). The analysis consisted in recording the % tail DNA on 50 cells per gel using the Comet IV software (Perceptive Instruments, Suffolk, UK). Experiments were performed three times independently, with two technical replicates per condition in each independent experiment. The cells exposed for 24 h to 100 µM of methyl methanesulfonate were used as positive control, and one control slide exposed to H_2_O_2_ was used as positive control for the electrophoretic migration.

### 2.9. CD11b membrane labelling

Cells previously exposed to pigments for 24h are then labeled with anti-CD11b antibody (BD Pharmigen, catalog number 553310, Becton Dickinson Biosciences, Le Pont-de-Claix, France) as follows: in a 96-well round-bottom plate, 200 µL of cell suspension containing 1 million cells/well is added to each well. The plate is then centrifuged for 5 minutes at 800 g. After turning the plate upside down, 50 µL of Fc block diluted at 10 µg/mL in PBS+2% SVF were added to each well. The plate was then left to incubate for 10 minutes at room temperature. Then 150 µL of PBS+2%SVF were added and centrifuged at 800 g. After turning the plate upside down, 50µL of CD11b antibody at 20µg/mL were added. The plate was then incubated for 20 minutes in the dark and at room temperature. After incubation, the cells were rinsed by adding 150 µL of PBS+2% SVF to each well. The plate was centrifuged for 5 minutes at 800 g and then emptied by turning the plate upside down. Finally, the cells were resuspended in PBS + 2% SVF + propidium iodide (1 µg/mL final concentration) and transferred to FACS tubes (final volume = 300 µL).

### 2.10. Numerical analysis

A Student’s *t*-test was applied to all the results. Data are presented as mean +/- standard deviations; *p<0.05, **p<0.01, ***p<0.001. The systematic use of viability markers (propidium iodide or Sytox red) allows for analysis of live cells only. Biological experiments were performed on four independent biological replicates.

## 3. Results

The first step of the study focused on the pigments (cobalt-and zinc-compounds) themselves. The physico-chemical characteristics of the pigments were first evaluated. Then their uptake at non-toxic doses (Lethal Dose 20) by macrophages and their possible dissolution were assessed. In a second step, we focused on the physiological and functional effects of macrophages resulting from the internalization of these pigments.

We chose to compare the effects of pigments on J774 macrophages and present the results corresponding to the two schemes of exposure tested. The acute effects of cobalt- and zinc-based pigments were compared to recovery exposure in order to visualize specific persistent effects. For the two schemes of exposures, a corundum control was used in order to distinguish the effects due to the internalization of particles from the specificities of the pigmentary materials tested.

### 3.1. Characterization of the pigments used

#### 3.1.1. Physical characterization of pigments via Dynamic Light Scattering and Transmission Electron Microscopy

The size (hydrodynamic radius) and the size distribution (polydispersity percentage) of the pigments in suspension were characterized using Dynamic Light Scattering (DLS). Size distribution is deemed monodisperse when the polydispersity percentage is inferior to 15%. The results in Figure 1A and supplementary Data 1 show a high diversity of sizes for all the pigments tested. Consequently, dispersity cannot be measured. However, the average hydrodynamic radius of each pigment tested was very different. Indeed, the average hydrodynamic radius of PV14 was approximately 994 nm in H2O medium and 860 nm in DMEM culture medium. For PG19, the radius size was 196 nm in H2O and 98 nm in DMEM. For PB28, the radius size was 517 nm in H2O and 441 nm in DMEM. Lastly, for PW4, the radius size was 242 nm in H2O and 177 nm in DMEM. Size variation between DMEM medium and H2O was not significant. Transmission Electron microscopy (TEM) (Figure 1B) confirmed a high diversity of sizes and shapes for all particle suspensions, both pigments and corundum. TEM microscopy revealed the non-negligible presence of nanoparticles with a size of about 50 nm in the case of PG19. Microscopic observations also showed the presence of nanostructured elements with at least 1 dimension in nanometer size. This can be easily observed in the cases of PB28 and PW4. Supplementary data 1 provides more precise measurements (not only the average size) recorded via DLS and shows the presence of submicrometer objects for all the pigments tested. For example, in the case of PB28, some objects have a hydrodynamic radius between 70 and 113 nm. In the cases of PV14 and PG19, we observed objects with a hydrodynamic radius between 2 and 20 nm, and between 8 and 110 nm, respectively. Finally, in the case of PW4, the objects observed have a radius between 6 and 124 nm. The results confirm the presence of nano-elements and sub-micrometric particles in addition to microparticles in the pigments tested.

#### 3.1.2. Chemical characterization of pigments after their dissolution in acidic medium via Inductively Coupled Plasma atomic emission spectroscopy

For three of the pigments used in this study, namely Pigment Violet 14, Pigment Blue 28 and Pigment White 4, the formulas are known (cobalt phosphate Co_3_(PO4)_2_, CoAl_2_O_4_ and ZnO, respectively). Pigment Green 19 is described as a mixture of cobalt and zinc oxide (Rinmann’s green) in undisclosed proportions, so that the chemical composition of the pigment had to be determined. To this end, we analyzed the three cobalt-based-pigments to determine their content of aluminum, zinc, and cobalt. For the purpose of comparison, corundum (Al_2_O_3_ spinel) was included in the analysis, as it has a similar spinel structure as Pigment Blue 28. Pigment White 4, which is also composed of zinc oxide, was analyzed in order to be compared with Pigment Green 19.

The results, shown in Table 1, allow to draw several conclusions.

**Table 1:** metal analysis of the pigments.

|  | Al (µg/l)<br>Measured after mineralization | Co (µg/l)<br>Measured after mineralization | Zn (µg/l)<br>Measured after mineralization |
| --- | --- | --- | --- |
| Negative control | 82.9 | 0 | 156 |
| Arabic gum 500 µg/ml | 158 | 0 | 77.4 |
| Corundum 2.5 µg/ml | 90 | 0 | 64.5 |
| Corundum 500µg/ml | 448 | 0 | 129 |
| PV14 2.5 µg/ml | 92.9 | 798 | 76.6 |
| PG19 2.5 µg/ml | 139 | 48 | 1810 |
| PB28 500 µg/ml | 2600 | 641 | 85.2 |
| PW4 2.5 µg/ml | 57.2 | 42.7 | 1420 |

As the input concentrations of the pigments are known, these experimental results allowed to have a clearer understanding of the mineralization efficiency for each pigment (the experiment was performed only once). First, the mineralization/measurement efficiency of PV14 was around 60%. Second, the mineralization/measurement efficiency of both spinels (corundum and PB28) was very low, below 1%.

Third, the mineralization/measurement efficiency of PW4 was 70%. Finally, the mineralization/measurement efficiency of PG19 was close to quantitative (92% yield) and allowed to determine the relative proportion of zinc and cobalt in the pigment. Thus, the pigment used in this study could be described as Co_0.028_Zn_0.912_O.

### 3.2. Determination of pigment dose for cellular assays

To carry out the following functional tests, it was necessary to determine the concentration of pigments allowing to obtain the best compromise between a good viability and observation of biological effects. We thus decided to limit the concentration at or just below the LD 20, in order to avoid excess mortality in case of slight variations of the applied doses. Given the particulate nature of the pigments, such variations are likely to occur. Because nanoparticles are known to interfere with many methods of viability estimation [64], we chose a dye exclusion test. According to the data displayed in Figure 3, we considered that the optimal dose was 2.5 µg/ml for PV14, PG19 and PW4, and 500 µg/ml for PB28.

**Figure 3.**
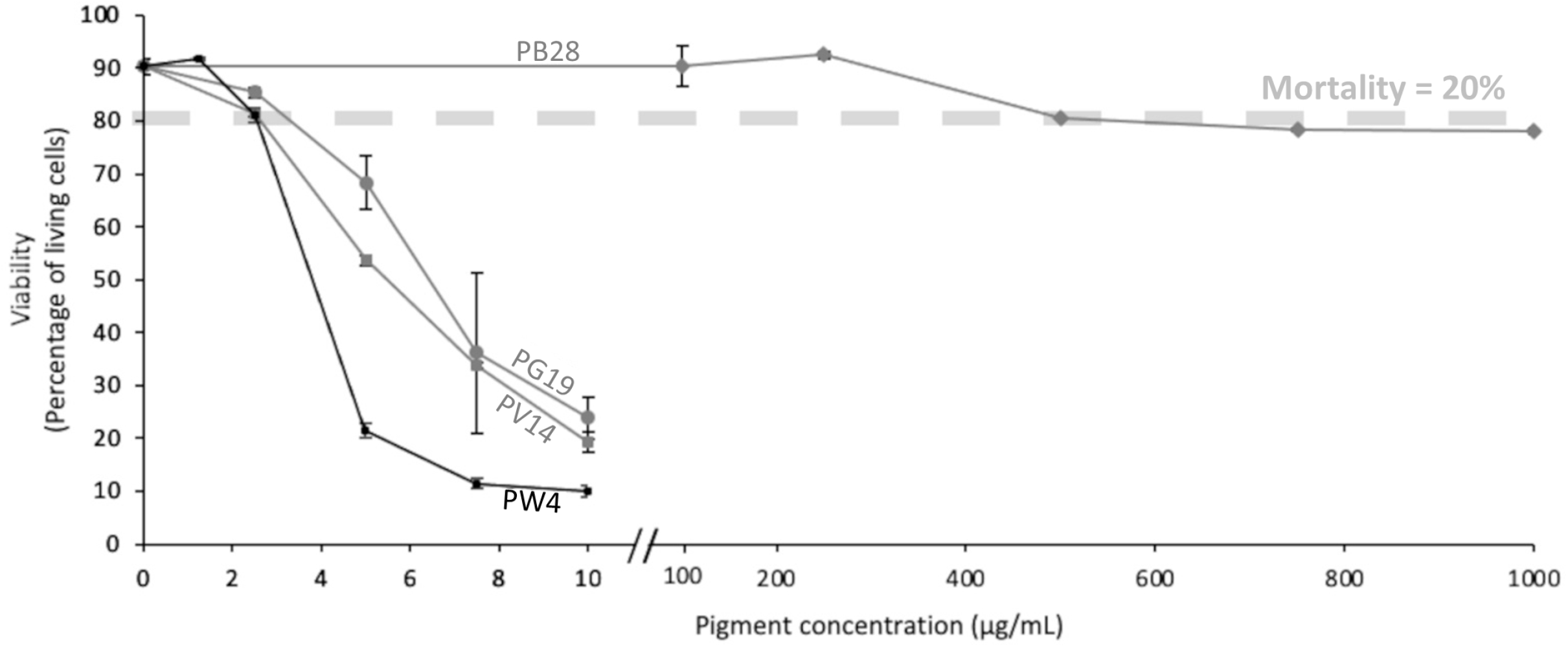
Viability of J774 cells. Cells were exposed to variable doses of pigments (PV14, PG19, PW4 and PB28) for 24h. Viability was measured using propidium iodide (1µg/ml).

### 3.3. Confirmation of the uptake and measurement of the dissolution rate of pigments into macrophages

#### 3.3.1. The pigments are internalized by macrophages

The pigments chosen for this study were made fluorescent in order to visualize at very low concentrations the presence of pigments inside the cells. Figures 4A and 4B obtained by confocal microscopy show the presence of pigments inside the macrophages proving that they are really internalized by the macrophages and not only deposited on the surface of the cells membranes. The presence of pigments was visualized in the cytosol and not in the nucleus. At 2.5 µg/mL conditions, only corundum, PG19 and PW4 were able to made fluorescent. The protocol did not allow successful covalency of rhodamine isothiocyanate with PV14. For higher pigment concentrations (Corundum 500 µg/mL, and PB28 500 µg/mL), better visualization of their presence in large quantity inside the cells is shown in supplementary data 2.

**Figure 4.**
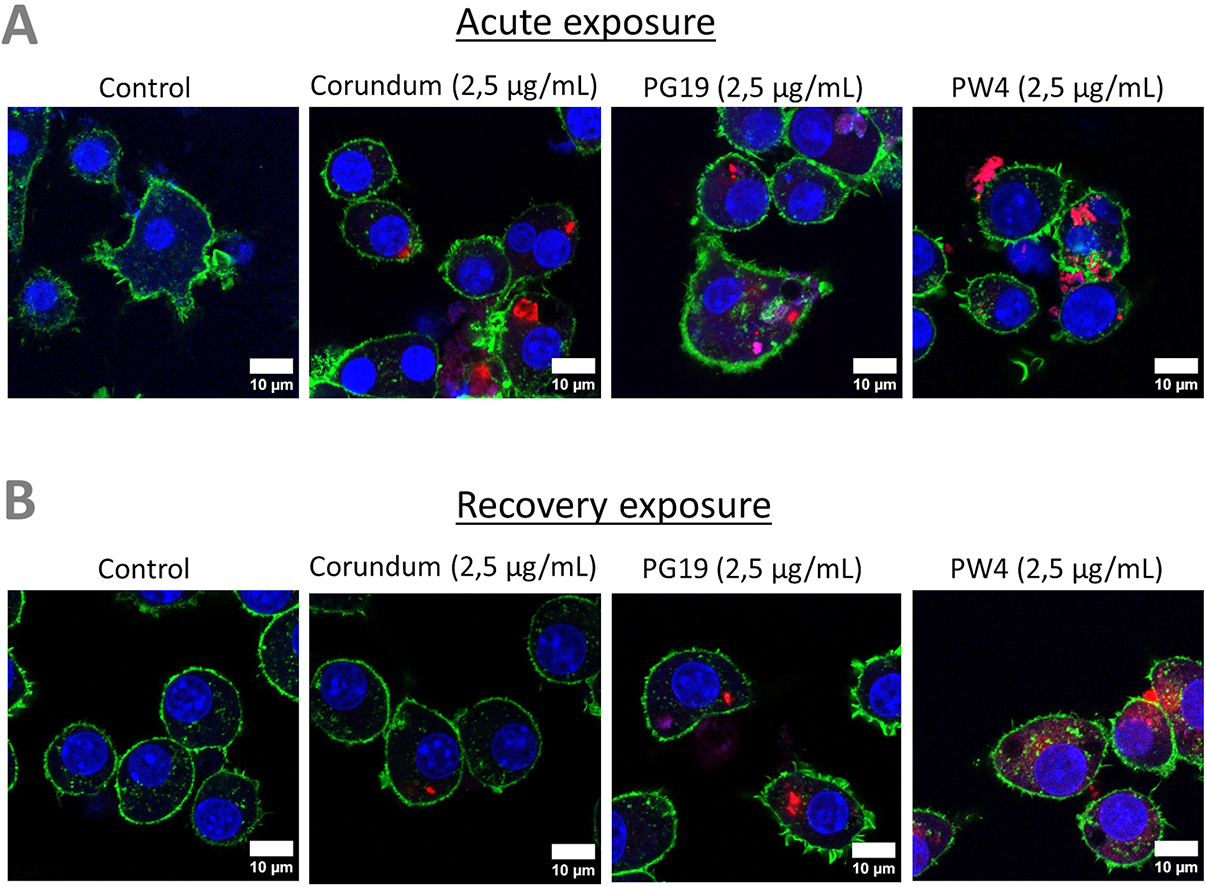
Visualization of pigment uptake with Rhodamine isothiocyanate (RITC)-coupled pigments. Cells were exposed to 2.5 µg/mL of fluorescent corundum or pigments (PG19 and PW4) for 24h. Green = actin dyed with Phalloidin-Atto 390; blue = nucleus dyed with Sytox^TM^ Red Dead Cell Stain; red = RITC-pigment. Scale bare = 10 µm.

#### 3.3.2. The pigments are slowly degraded in macrophages

Macrophages engulf particles via the phagolysosomal pathway. Consequently, pigments internalized by macrophages were certainly exposed to an environment that is both acidic and oxidizing, which could cause metal ions to be released from the pigments into the cells. To evaluate this phenomenon, we measured the soluble ion content of cellular extracts after exposure to the pigments, either at the end of the 24h exposure phase or at the end of the 72h post-exposure period. The results, displayed in Table 2, show distinct differences between the end of the exposure period on the one hand, and the end of the recovery period on the other.

**Table 2:** intracellular ion content after exposure to pigments.

|  |  | Acute |  |  | Recovery |  |  |
| --- | --- | --- | --- | --- | --- | --- | --- |
|  |  | ng Al/mg cell proteins | ng Co/mg cell proteins | ng Zn/mg cell proteins | ng Al/mg cell proteins | ng Co/mg cell proteins | ng Zn/mg cell proteins |
| Neg ctrl |  | 166±117 | 18.6±20.9 | 11.1±17.1 | 16.9±41.4 | 15.2±14.2 | 59.5±19.3 |
| Arabic gum | 500 µg/ml | 29.5±35.5 | 40.8±46.7 | 43.2±38 | 29.8±30.2 | 13.2±14.9 | 12.5±21.5 |
| Corundum | 2.5µg/ml | 117.3±48.2 | 55.7±56.5 | 22.9±12.2 | 0.38±0.94 | 56.2±32.2 | 29.9±26.5 |
| Corundum | 500 µg/ml | 748.7±201.7* | 17.7±15.7 | 34.1±30.9 | 116.3±103.9 | 59.7±72.8 | 26±23.9 |
| PV14 | 2.5µg/ml | 17.7±24.7 | 225.2±118.5* | 17.4±14.3 | 0 | 53.7±41.7 | 34.8±26.8 |
| PG19 | 2.5µg/ml | 35.7±34.1 | 42.9±36.5 | 70.7±16* | 30.8±35 | 25.6±14.7 | 34.8±14.9 |
| PW4 | 2.5µg/ml | 25.4±34.4 | 34±27.4 | 97.1±24 | 60±69.2 | 6.3±7.1 | 38.6±12.2 |
| PB28 | 500 µg/ml | 1083.6±181.9* | 125.15±47.2* | 56.8±58.6* | 477.3±295.3* | 145.7±59.8* | 34.1±38.2 |

\* Significantly different from control in two-tailed Mann-Whitney U test

At the end of the exposure period, a significant ion release was detected for PV 14 and PB 28. However, this release must be put in perspective with the total metal input. For example, the total input of 1 mg PB28 corresponds to 330 µg of cobalt, of which only 28 ng are detected as intracellular and soluble, i.e. a 0.01% solubilization. Furthermore, the amounts of aluminum and cobalt released from PB28 should be very similar, which is not the case in our data. This shows that cells do not accumulate all ions passively, but may either solubilize ions differently or excrete some ions more than others. PV14 appears to be more soluble, with 43 ng of cobalt detected as intracellular and soluble, to be compared with the 2.4 µg of cobalt input, i.e. a 1.8% solubilization.

The results are more difficult to interpret at the end of the recovery period, because the number of cells can vary from one condition to another, for example because some cells detach from the plate, as indicated by lower protein contents. However, a sustained release of cobalt can be seen from PV14 and from PB28.

This soluble ionic release in the cells may therefore play a role in the physiology and functionality of macrophages, which was evaluated next.

### 3.4. From disruption of antioxidant defenses to DNA damage: an effect related to the type of pigment and its concentration

#### 3.4.1. Intracellular glutathione levels are weakly altered upon pigment ingestion

One of the main detoxification mechanisms triggered by cells against xenobiotics (such as metals) is glutathione (GSH). This intracellular molecule has antioxidant properties and reduces the production of reactive oxygen species (ROS). In this experiment, we first identified three possible types of cell populations: GSH “high” intensity cells similar to the control without treatment, GSH “intermediate” intensity and GSH “low” intensity (see examples in Supplementary data 3). The percentage measured of these different populations according to pigments is presented in Figure 5A (for percentage of GSH “High” intensity), Figure 5B (for GSH “intermediate” intensity and Figure 5C (for GSH “low” intensity). Results show a very slight (but significant) decrease in the number (%) of “high” GSH cells for PG19 (2.5 µg/mL) and PW4 (2.5 µg/mL) only in acute exposure. These results are confirmed when looking at the “intermediate” GSH population and they only confirmed for PW4 for the percentage of “low” GSH population. On the other hand, for PV14 (2.5 µg/mL), this small but significant decrease persists during recovery exposures. This trend is validated by the emergence of a “low” GSH population (3.7 %) in recovery condition. For the PB28 condition, a slight decrease in the number of “high” GSH cells and the appearance of an “intermediate” (12%) and “low” (2.5%) intensity population were observed in acute exposure. However, a more significant effect was visible in recovery exposure. Indeed, only 46 % of cells are “high” GSH intensity and about 50 % of cells became “low” GSH with a small percentage (4%) of “intermediate” GSH. These results are significant in comparison to the condition without treatment and Corundum 2,5 µg/mL or 500 µg/mL. The weak overload observed for corundum 500 µg/mL seems to be temporary, indeed this effect is not found in recovery exposure. The positive control (in the presence of 50 µM Menadione) shows a disappearance of 80 to 85 % of the GSH ‘High” population and the appearance of an “intermediate” population in the majority.

**Figure 5.**
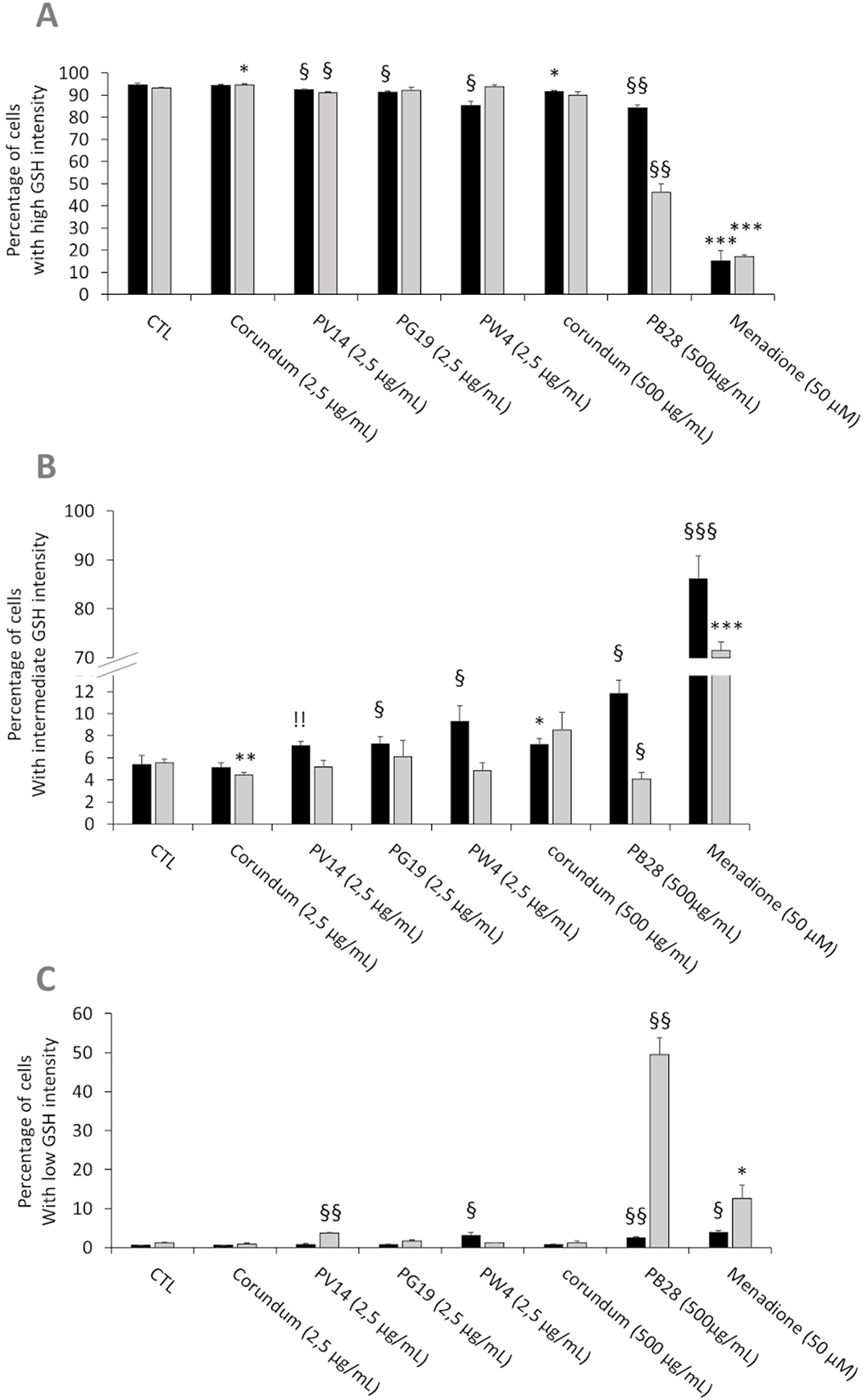
Intracellular glutathione measurement. **Black bars: acute exposure (24 hours). Grey bars: exposure for 24 hours followed by a 3-day post-exposure recovery period**. Panel A- Percentage of cells with high intensity glutathione. Panel B- Percentage of cells with intermediate intensity glutathione. Panel C- Percentage of cells with low intensity glutathione. Menadione is used as positive control. Significance of symbols: *: different from negative control only; §: different from both negative and corundum controls 500 µg/ml. Number of symbols: 1: p<0.05; 2: p<0.01.

#### 3.4.2. Pigments induce a small increase in Reactive Oxygen Species levels, and show weak genotoxicity

The accumulation of reactive oxygen species (ROS) in the cell can have cytotoxic and genotoxic effects leading to functional perturbations. Thus, the accumulation of ROS and the search for DNA damage were carried out. Experiments using the DHR123 method showed (Figure 6A) a significant increase in ROS only for the PB28 condition (500 µg/mL) compared to its untreated and corundum (500 µg/mL) controls. This increase is visible in acute exposure and is maintained during recovery exposure. As there is no effect in the presence of corundum (500 µg/mL), the increase of ROS was specific to the PB28 pigment (500 µg/mL). The other pigments did not seem to increase the quantity of intracellular ROS compared to the control without treatment. However, a very slight significant effect was visible in comparison with the corundum condition (2.5 µg/mL). This significance can be explained by an almost inexistent standard deviation for this control. The positive control in the presence of menadione 50 µM showed a strong increase of ROS which is multiplied by 4 and 3 compared to the PB28 condition (500 µg/mL) in acute or recovery exposure.

**Figure 6.**
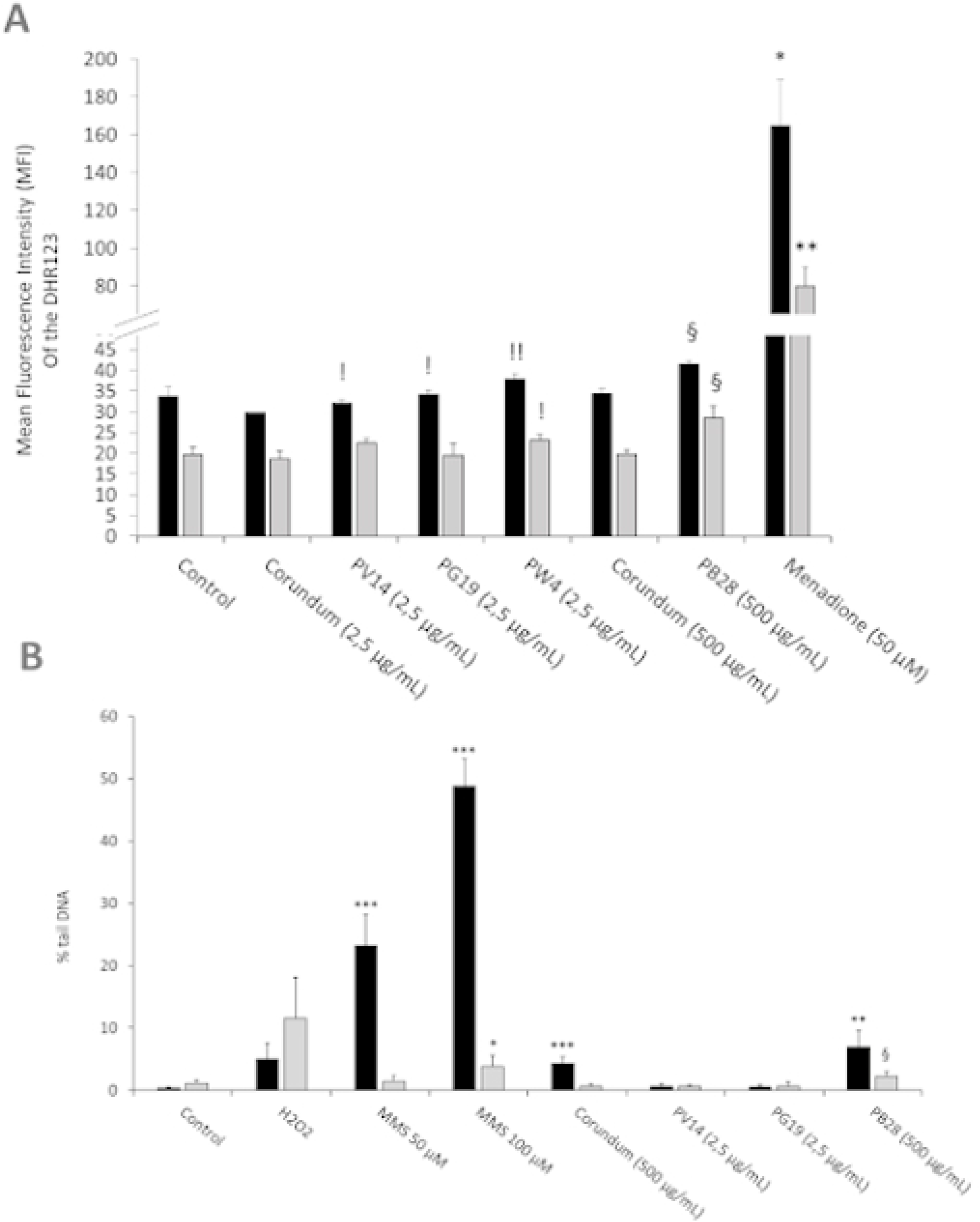
ROS and Genotoxicity assessment. Panel A- Measurement of intracellular ROS with DHR123 (500ng/mL). Menadione is used as positive control. Panel B- DNA damage determined by a comet assay. The percentage of DNA in the comet tail is measured as an indicator of the extent of damage to DNA. Methyl methanesulfonate (MMS) and hydrogen peroxide are used as positive controls. **Black bars: acute exposure (24 hours). Grey bars: exposure for 24 hours followed by a 3-day post-exposure recovery period**. Significance of symbols: *: different from negative control only; §: different from both negative and corundum controls (500 µg/ml). !: different from corundum control only. Number of symbols: 1: p<0.05; 2: p<0.01; 3: p<0.001.

To follow up on the ROS results obtained previously and because of cobalt is known to be genotoxic [28], we investigated the possible genotoxic effects and tested it via a comet assay. The results presented in Figure 6B showed that neither corundum nor the pigments had any affect at low concentrations (2.5µg/ml), whether immediately after exposure or after the recovery period. However, a significant increase in DNA damage was observed immediately after exposure to high concentrations of particles (either corundum or PB28). The effect disappeared at the end of the recovery period in the case of corundum, but not in the case of PB28 where a sustained effect was detected, even though it lost intensity after recovery. PW4 was not tested because the genotoxicity of zinc oxide on macrophages has already been documented [50, 65]. The result showed that the concentration ≤ 3.5 or 4 µg/mL in ZNO nanoparticles did not induce DNA damage to macrophages.

The data shown in supplementary data 4 on a mitotic cell showed on the one hand the direct contact of pigment particles with chromosomes and on the other hand that the particle distribution within the cell was not symmetrical compared to the cell division plane. This suggested that the particle concentration may be different between daughter cells [66] and that the cell division process itself might be disturbed by the presence of high concentrations of particles.

### 3.5. Pigments induce a small decrease in phagocytic activity, and do not damage lysosomes

#### 3.5.1. Measurement of phagocytic activity

To determine whether the pigments induce changes to the main functions of macrophages, a phagocytosis assay was carried out for acute and recovery exposure. The results in Figures 7A and 7C showed that under acute exposure, PV14 (2.5 µg/ml), PG19 (2.5 µg/mL) and PW4 (2.5 µg/mL) induced a decrease in the proportion of phagocytic cells in comparison to control without treatment and corundum (2.5 µg/ml) control. For PV14 (2.5 µg/mL), the percentage of phagocytic cells decreased by about 10% in comparison to the two controls. For PG19 (2.5 µg/ml), we observed a decline of 14 and 13% in comparison to control without treatment and corundum (2.5 µg/ml), respectively. For PW4, there was a decrease of 12 and 11% in comparison to control without treatment and corundum (2.5 µg/ml), respectively. Under recovery exposure, the proportion of phagocytic cells remained significantly lower than controls for PG19 (2.5 µg/mL) and PW4 (2.5 µg/mL) only. Indeed, for PG19, the percentage of phagocytic cells decreased by 18% and 14% in comparison to control without treatment and corundum (2.5 µg/mL). The decline observed for PW4 was similar (between 19 and 15%). PV14 (2.5 µg/mL) did not induce significant changes in comparison to control without treatment. Thus, at 2.5 µg/mL, short-term and persistent effects on the phagocytic activity were only observed for PG19 (2.5 µg/mL) and PW4 (2.5 µg/mL). These two pigments contain zinc (in addition to cobalt, for PG19). PV14 (2.5 µg/mL) is the only pigment that induced solely an acute effect. Under acute exposure, the results presented in Figures 7B and 7D showed that PB28 (500 µg/mL) and the corundum control (500 µg/ml) induced a significant decrease in phagocytic cells (−22% and −28%, respectively) in comparison to control without treatment. No significant differences between the corundum (500 µg/ml) and PB28 (500 µg/ml) conditions were observed. Therefore, the reduction of the phagocytic activity was due to a high concentration of particles, regardless of the nature of the particulate material. However, these decreases were maintained (and, perhaps, aggravated) only for PB28 (500 µg/ml) under recovery exposure. Indeed, the percentage of phagocytic cells declined by 30%.

**Figure 7.**
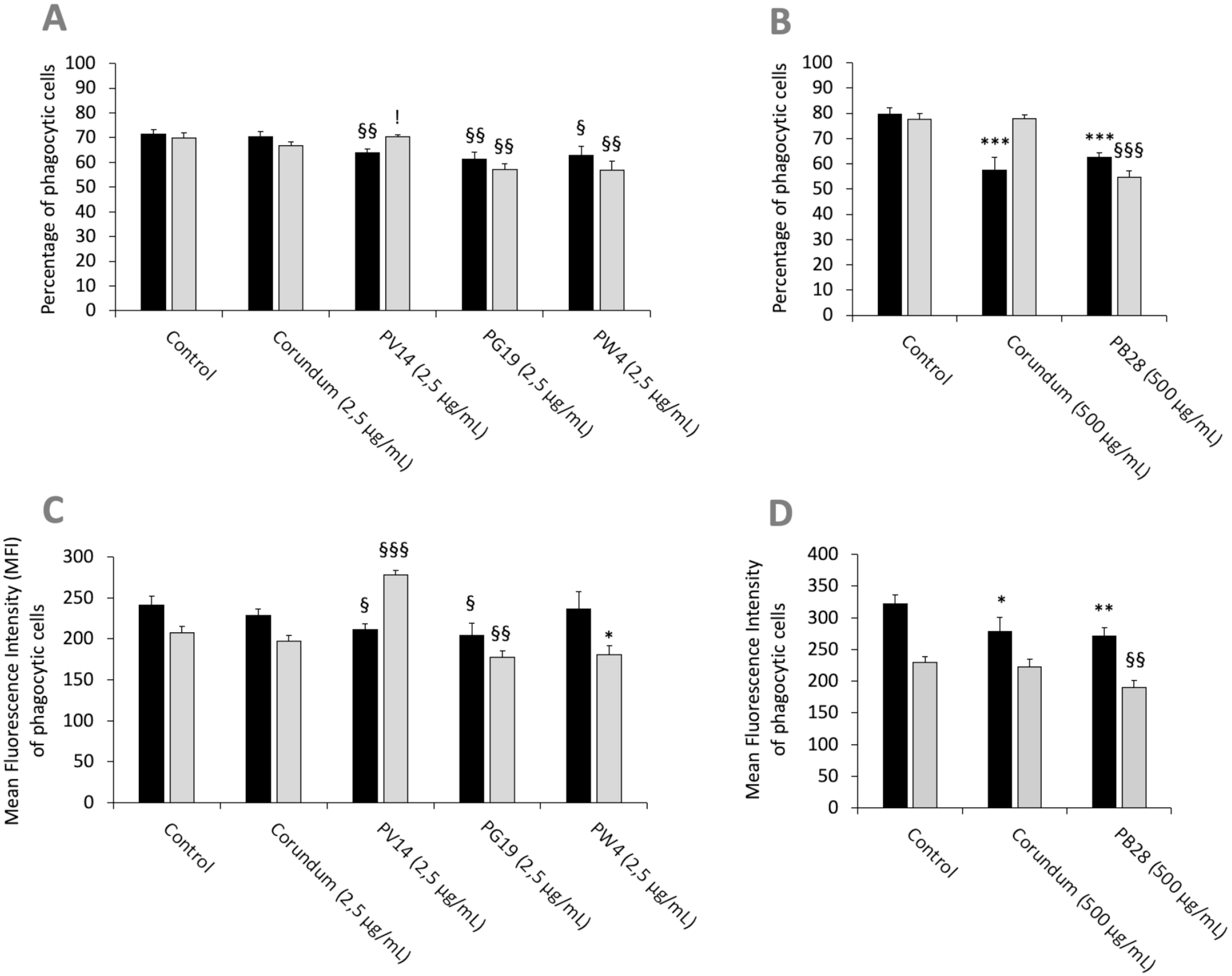
Phagocytic capacity. **Black bars: acute exposure (24 hours). Grey bars: Exposure for 24 hours followed by a 3-day post-exposure recovery period.** Panel A: conditions with 2.5 µg/ml of particle suspension. Percentage of cells able to phagocytose fluorescent FITC-labeled latex beads (positive cells). Panel B: conditions with 500 µg/ml of particle suspension. Percentage of cells able to phagocytose fluorescent FITC-labeled latex beads (positive cells). Panel C: measurement of the phagocytic ability (mean fluorescent intensity) of positive cells. Conditions with 2.5 µg/ml of particle suspension. Panel D: measurement of the phagocytic ability (mean fluorescent intensity) of positive cells. Conditions with 500 µg/ml of particle suspension. Significance of symbols: *: different from negative control only; §: different from both negative and corundum controls; !: different from corundum control only. Number of symbols: 1: p<0.05; 2: p<0.01; 3: p<0.001.

Under acute exposure, we observed a significant decrease in the Mean Fluorescence Intensity (MFI) for PV14 (2.5 µg/mL) (MFI = 211) and PG19 (2.5 µg/mL) (MFI = 205) in comparison to their controls (MFI = 242 for control without treatment and MFI = 229 for corundum (2.5 µg/mL)). Under recovery exposure, the decrease continued for PG19 (2.5 µg/mL) (MFI = 177) only. Surprisingly, for PV14, the MFI significantly increased to 278.

For PB28, we observed a similar decrease in the MFI compared to corundum (500 µg/mL) under acute exposure (MFI = 279 and 271, respectively, instead of 321 for control without treatment). Under recovery exposure, the decline of the MFI for PB28 continued in comparison to controls.

#### 3.5.2 Measurement of lysosomal activity

Besides the phagocytic activity, we tested whether the pigments altered the lysosomal activity. The neutral red uptake assay allows a crude evaluation of the endosomal and lysosomal activities as it requires an active proton pump and undamaged lysosomes. The results in Figures 8A and 8B showed no significant differences regarding the uptake of neutral red between the conditions with and without pigment, under both acute and recovery exposure. Thus, no damage to the endosome-lysosome pathway was observed, thereby confirming that the decrease in phagocytic ability in the presence of PB28 (500 µg/ml) and corundum (500 µg/ml) impacted solely the phagocytosis process without modifying lysosomal integrity.

**Figure 8.**
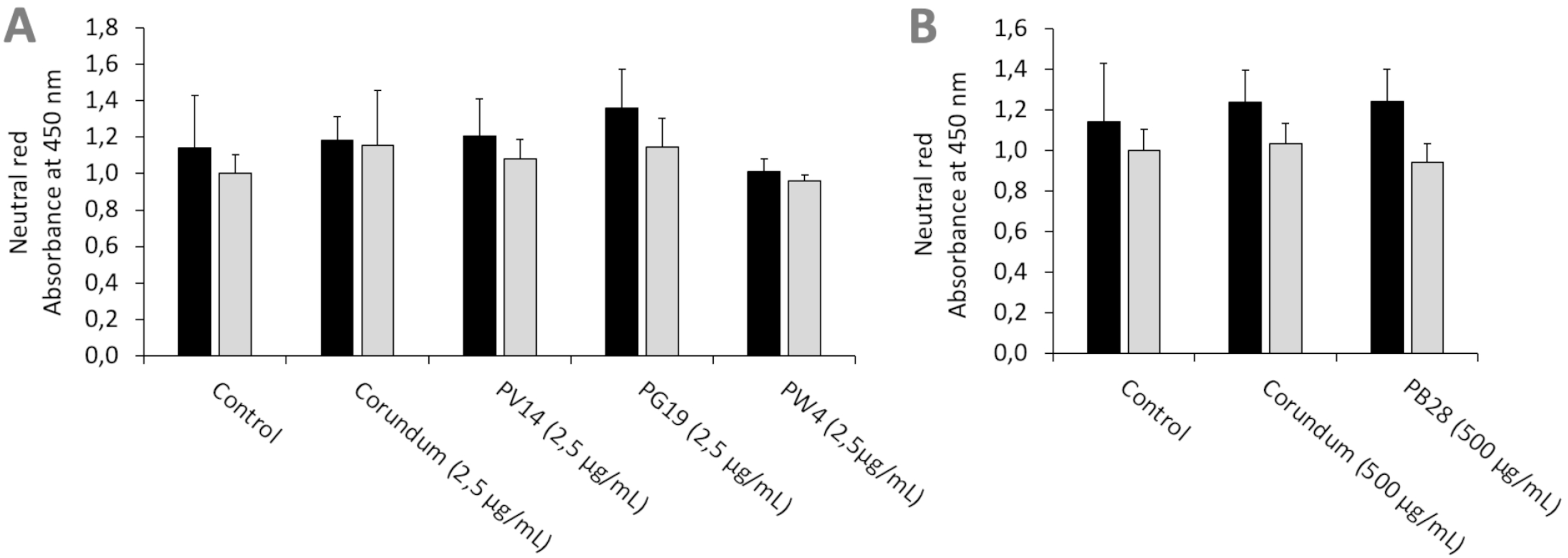
Neutral red uptake. **Black bars: acute exposure (24 hours). Grey bars: exposure for 24hours followed by a 3-day post-exposure recovery period.** Panel A: conditions with 2.5 µg/ml of particle suspension. Panel B: conditions with 500 µg/ml of particle suspension. After elution, the absorbance of the red color released was read at 540 nm.

In order to validate that the phagocytosis results obtained by flow cytometry are not due to an adsorption phenomenon, observations (Supplementary data 2) via confocal microscopy validated the presence of the beads (used in classical phagocytosis assays) inside the cells and not only on their surface membrane. Interestingly, for PB28 (500 µg/mL) phagocytosis seems to stop just after that beads penetrated the cell membrane due to a high density of pigments. This phenomenon is only showed in recovery exposure. The density of pigment seems to be lower in acute exposure.

### 3.6. Inflammatory response

The inflammatory response of macrophages was tested with nitric oxide (NO) and cytokine assays. The responses, under all conditions, were assessed in the presence or absence of inflammatory stimuli such as LPS.

#### 3.6.1. Pigments alter nitric oxide production

We first tested the intrinsic effects of pigments without LPS stimulation. In acute exposure mode, the results in Figures 9A and 9B show that the production of NO increased strongly and significantly for PV14 (2.5 µg/ml), i.e. 9.5 µM in comparison to 4.4 µM for control without treatment and 2.8 µM for corundum (2.5 µg/mL) control. For PG19 (2.5 µg/mL) and PW4 (2.5 µg/mL), a less significant increase was also observed. Indeed, the secretion of NO was 6.4 µM for PG19 (2.5 µg/mL) and 5.2 µM for PW4 (2.5 g/mL). Thus, this pro-inflammatory effect was specific to these pigments since no increase in NO secretion was observed for corundum (2.5 µg/mL). Moreover, PB28 (500 µg/mL) did not induce an intrinsic increase in NO secretion despite a high concentration.

**Figure 9.**
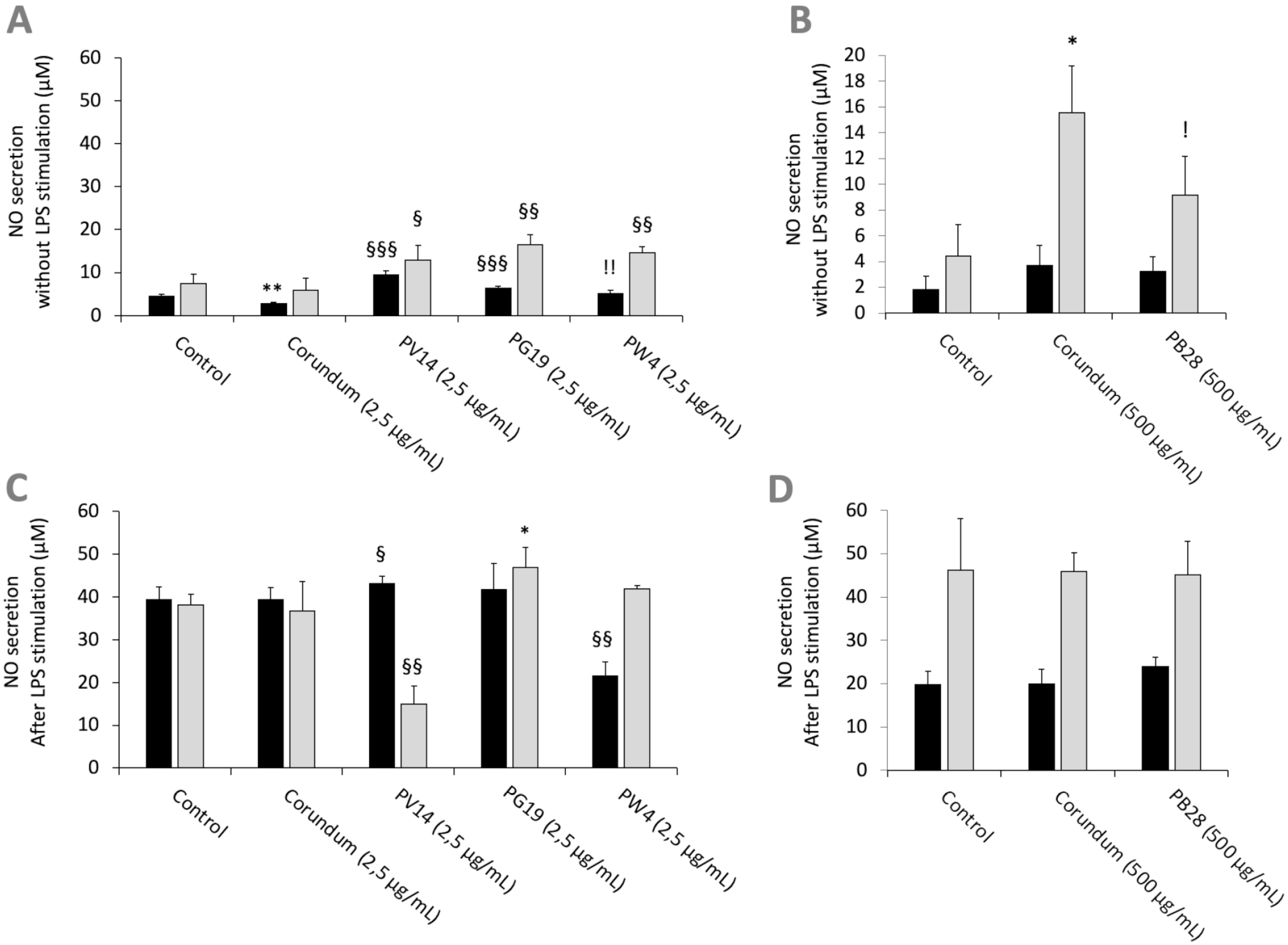
NO secretion. **Black bars: acute exposure (24 hours). Grey bars: exposure for 24 hours followed by a 3-day post-exposure recovery period.** Panel A: NO secretion without LPS stimulation. Conditions with 2.5 µg/ml of particle suspension. Panel B: NO secretion without LPS stimulation. Conditions with 500 µg/ml of particle suspension. Panel C: NO secretion with LPS stimulation. Conditions with 2.5 µg/ml of particle suspension. Panel D: NO secretion with LPS stimulation. Conditions with 500 µg/ml of particle suspension. Significance of symbols: *: different from negative control only; §: different from both negative and corundum controls; !: different from corundum control only. Number of symbols: 1: p<0.05; 2: p<0.01; 3: p<0.001.

Under recovery exposure, the results showed a persisting and significant increase in NO secretion in the presence of pigment in comparison to their controls. Indeed, NO secretion amounted to 12.8 µM for PV14 (2.5 µg/mL), 16.4 µM for PG19 (2.5 µg/mL) and 14.7 µM for PW4 (2.5 µg/mL), but only 7.4 µM for control without treatment and 5.8 µM for corundum (2.5 µg/mL). For corundum (500 µg/mL) and PB28 (500 µg/mL), the results were more difficult to interpret. Indeed, we observed a surprising increase in NO secretion in the presence of corundum (500 µg/mL) but not in the presence of PB28 (500 µg/mL). However, we can conclude that, contrary to others pigments with a smaller concentration, PB28 (500 µg/mL) cannot induce intrinsic effects on NO secretion after recovery exposure.

After stimulating cells with LPS for the last 18 hours of the experiment, in acute or recovery exposure modes, the results obtained with pigments in comparison to the condition without particles showed that the secretion of NO remained unchanged for cells exposed to corundum (2.5 µg/mL and 500 µg/mL), PB28 (500 µg/mL) and PG19 (2.5 µg/mL) (Figures 9C and 9D). Under acute exposure, we observed a slight trend toward an increase in the concentration of NO for PV14 (2.5 µg/mL) and a decrease in NO secretion (21.5 µM) for PW4 (2.5 µg/mL) in comparison to the two controls without treatment and corundum (2.5 µg/mL) (39.4 µM and 39.3 µM, respectively). Under recovery exposure, PG19 (2.5 µg/mL) and PW4 (2.5 µg/mL) did not induce any change to the secretion of NO in comparison to their controls. However, PV14 (2.5 µg/mL) induced a sharp decline (15 µM) in comparison to its controls i.e. 38.2 µM and 36.7 µM for control without treatment and corundum (2.5 µg/mL), respectively. Moreover, we observed that for PV14 (2.5 µg/mL-recovery), the level of NO was similar with or without LPS stimulation (15 µM with LPS stimulation and 12.8 µM without LPS stimulation). No significant changes were observed for PB28 (500 µg/mL) under recovery exposure despite a significant increase in NO secretion for corundum (500 µg/mL) in comparison to the control without treatment.

#### 3.6.2. Pigments induce a small pro-inflammatory effect, but alter cellular responses to bacterial components

In order to determine whether pigments induced an intrinsic inflammatory effect, as suggested by the nitric oxide results, the release of cytokines by cells was measured without LPS stimulation. In acute exposure mode, the results in Figures 10C and 10D revealed no changes to IL-6 secretion for the conditions PV14 (2.5 µg/mL), PG19 (2.5 µg/mL), and PW4 (2.5 µg/mL) in comparison to their controls. However, IL-6 concentration increased in the presence of corundum (500 µg/mL) and PB28 (500 µg/mL) in comparison to control without treatment while no significant differences were observed between PB28 (500 µg/mL) and corundum (500 µg/mL). Thus, this increase was due to the presence of a high concentration of particles, independently of the material.

**Figure 10.**
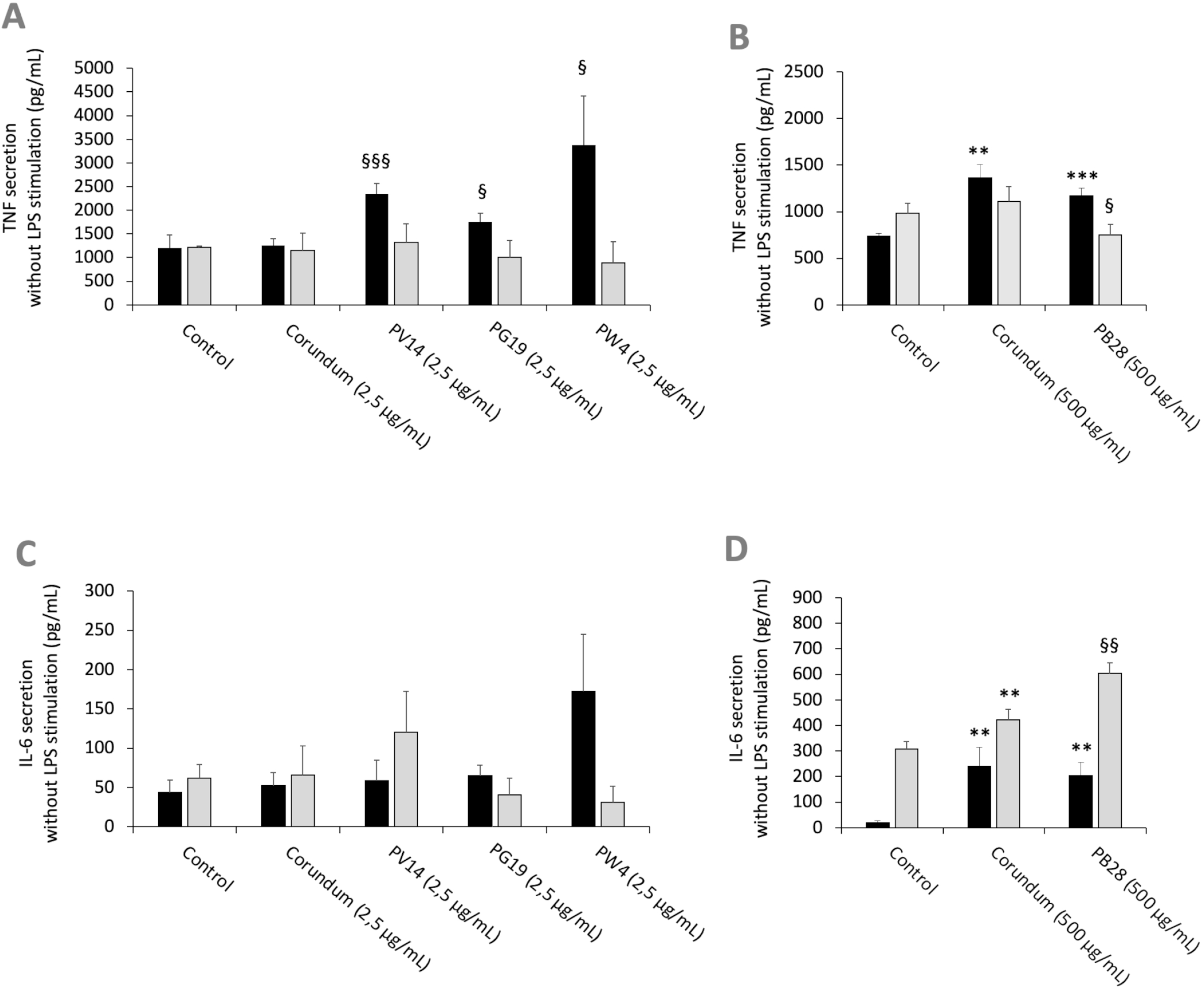
Cytokine secretion without LPS stimulation. **Black bars: acute exposure (24 hours). Grey bars: exposure for 24 hours followed by a 3-day post-exposure recovery period.** Panel A: TNF secretion. Conditions with 2.5 µg/ml of particle suspension. Panel B: TNF secretion. Conditions with 500 µg/ml of particle suspension. Panel C: IL-6 secretion. Conditions with 2.5 µg/ml of particle suspension. Panel D: IL-6 secretion. Condition with 500 µg/ml of particle suspension. Significance of symbols: *: different from negative control only; §: different from both negative and corundum controls. Number of symbols: 1: p<0.05; 2: p<0.01; 3: p<0.001.

For recovery exposure without LPS stimulation, no significant differences in IL-6 secretion were observed between the different conditions except for PB28 (500 µg/mL). In this case, IL-6 secretion increased significantly in comparison to its two controls. Part of the persisting effect observed was certainly due to the high concentration of particles because corundum (500 µg/mL) induced also a moderate increase in comparison to control without treatment.

The results in Figures 10A and 10B showed an increase in TNF secretion for PV14 (2.5 µg/mL), PG19 (2.5 µg/mL) and PW4 (2.5 µg/mL) in comparison to control without treatment and corundum (2.5 µg/mL). Both corundum (500 µg/mL) and PB28 (500 µg/mL) induced an increase in TNF in comparison to control without treatment. The results suggest that a high concentration of particles has an effect on TNF secretion independently of the material. Under recovery exposure, no significant differences were observed between the conditions tested except for PB28. Indeed, a slightly significant decrease in TNF concentration was observed: 752 pg/mL instead of 981 pg/mL and 1,111 pg/mL for control without treatment and corundum control (2.5 pg/mL), respectively.

After LPS stimulation and under acute exposure, IL-6 secretion (Figures 11C and 11D) remained unchanged for all the pigments tested in comparison to their controls. Conversely, under recovery exposure, the results showed a sharp drop in the secretion of IL-6 for PV14 (2.5µg/mL) (6079 pg/mL), PG19 (2.5 µg/mL) (4,874 pg/mL) and PW4 (2.5 µg/mL) (3,108 pg/mL) in comparison to control without treatment (12,334 pg/mL) and corundum control (2.5 µg/mL) (14,250 pg/mL). A significant decrease in IL-6 was also observed for PB28 (9,704 pg/mL) but only in comparison to control without treatment (22,870 pg/mL). Indeed, a slight decrease was also observed for corundum (500 µg/mL) (16,691 pg/mL). Thus, for PB28 (500 µg/mL), the decrease was mainly due to the high concentration of particles and was not material-dependent.

**Figure 11.**
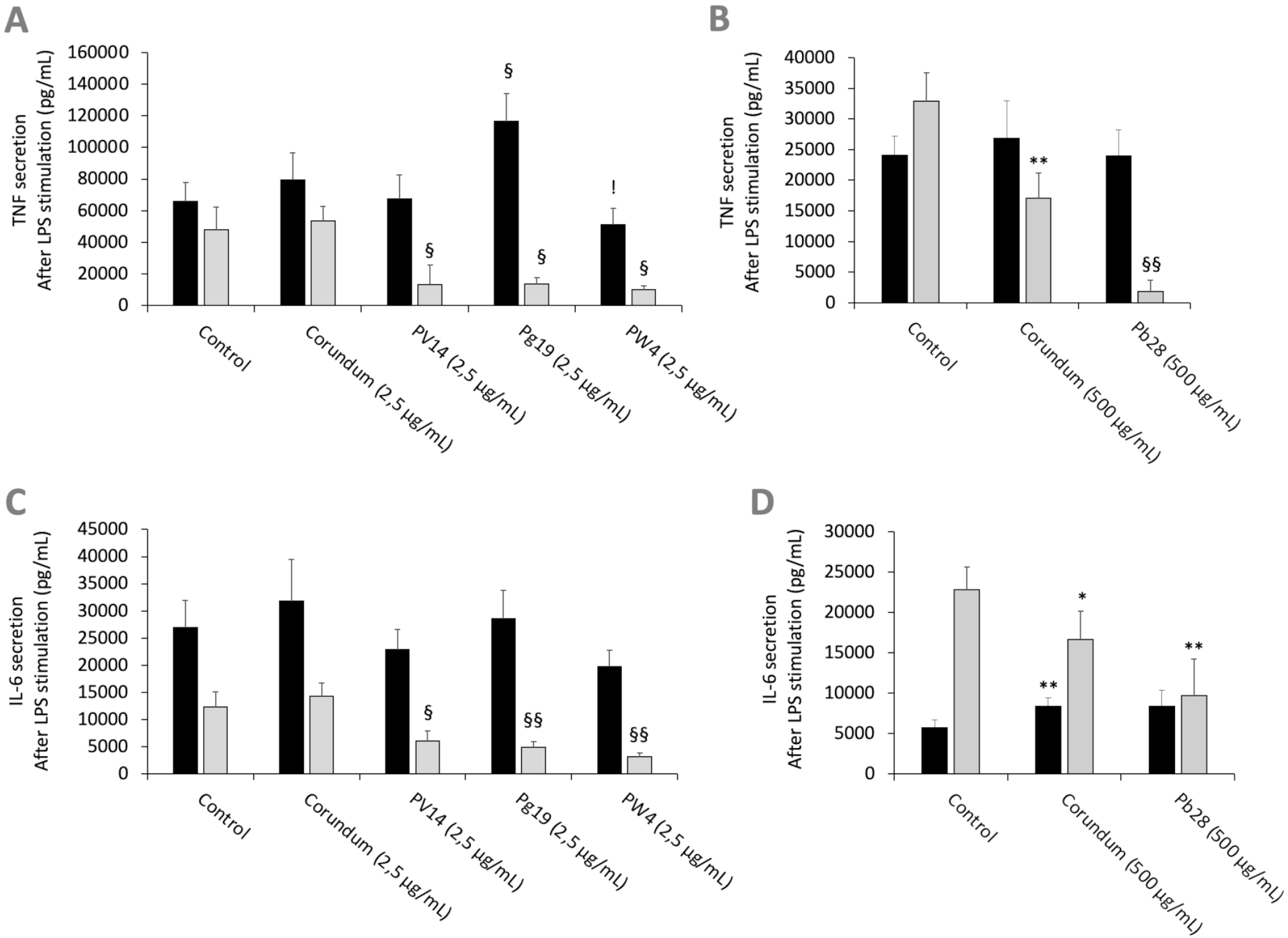
Cytokine secretion after LPS stimulation. **Black bars: acute exposure (24 hours). Grey bars: exposure for 24 hours followed by a 3-day post-exposure recovery period**. Panel A: TNF secretion. Conditions with 2.5 µg/ml of particle suspension. Panel B: TNF secretion. Conditions with 500 µg/ml of particle suspension. Panel C: IL-6 secretion. Conditions with 2.5 µg/ml of particle suspension. Panel D: IL-6 secretion. Condition with 500 µg/ml of particle suspension. Significance of symbols: *: different from negative control only; §: different from both negative and corundum controls; !: different from corundum control only. Number of symbols: 1: p<0.05; 2: p<0.01; 3: p<0.001.

The results in Figures 11A and 11B showed that under acute exposure and with LPS stimulation, no significant decrease in TNF secretion was observed for PV14 (2.5 µg/mL), PW4 (2.5 µg/mL), and PB28 (500 µg/mL). However, a significant increase was observed for PG19 (2.5 µg/mL) (11,6296 pg/mL instead of 65,810 pg/mL and 79,474 pg/mL for control without treatment and control corundum 2.5 pg/mL). Under recovery exposure, a sharp drop was observed (as in the case of IL-6 secretion). Indeed, for PV14, PG19, and PW4 (2.5 µg/mL), the concentration was 7,439 pg/mL, 13,887 pg/mL and 10,180 pg/mL, respectively, while for control without treatment and corundum (2.5 µg/mL), the concentration in TNF was 47,920 pg/mL and 53,374 pg/mL, respectively. In the case of PB28, the decline (1,796 pg/mL) was more drastic than in the case of the other pigments in comparison to controls without treatment (32,935 pg/mL) and corundum (500 µg/mL) (26,824 pg/mL).

In the case of PB28, the decrease was amplified by the high concentration of particles because a significant decrease was also observed in the case of corundum (500 µg/mL) in comparison to control without treatment.

#### 3.6.3. Pigments induce an alteration of the secretion of chemoattractants both in the absence an in the presence of bacterial components

In order to determine whether pigments induce an intrinsic effect on the production of chemoattractant molecules (in addition to pro-inflammatory cytokines) necessary for the recruitment of immune cells, the release of chemotactic cytokines MCP-1 and MIP-1α was measured without LPS stimulation.

In the absence of LPS stimulation, the results in Figure 12A and 12B showed a significant decrease in MCP-1 secretion for all pigments tested during acute exposure compared to their respective controls (No Treatment and Corundum). This decrease persisted significantly in recovery exposure for PV14 (2,5 µg/mL) and PB28 (500 µg/mL). However, for MIP-1α (Figure 12C and 12D) only PB28 (500 µg/mL) induced a significant decrease both in acute and recovery exposure in comparison to its controls. For all conditions without LPS stimulation, all values are between 2300 pg/mL and 4200 pg/mL

**Figure 12.**
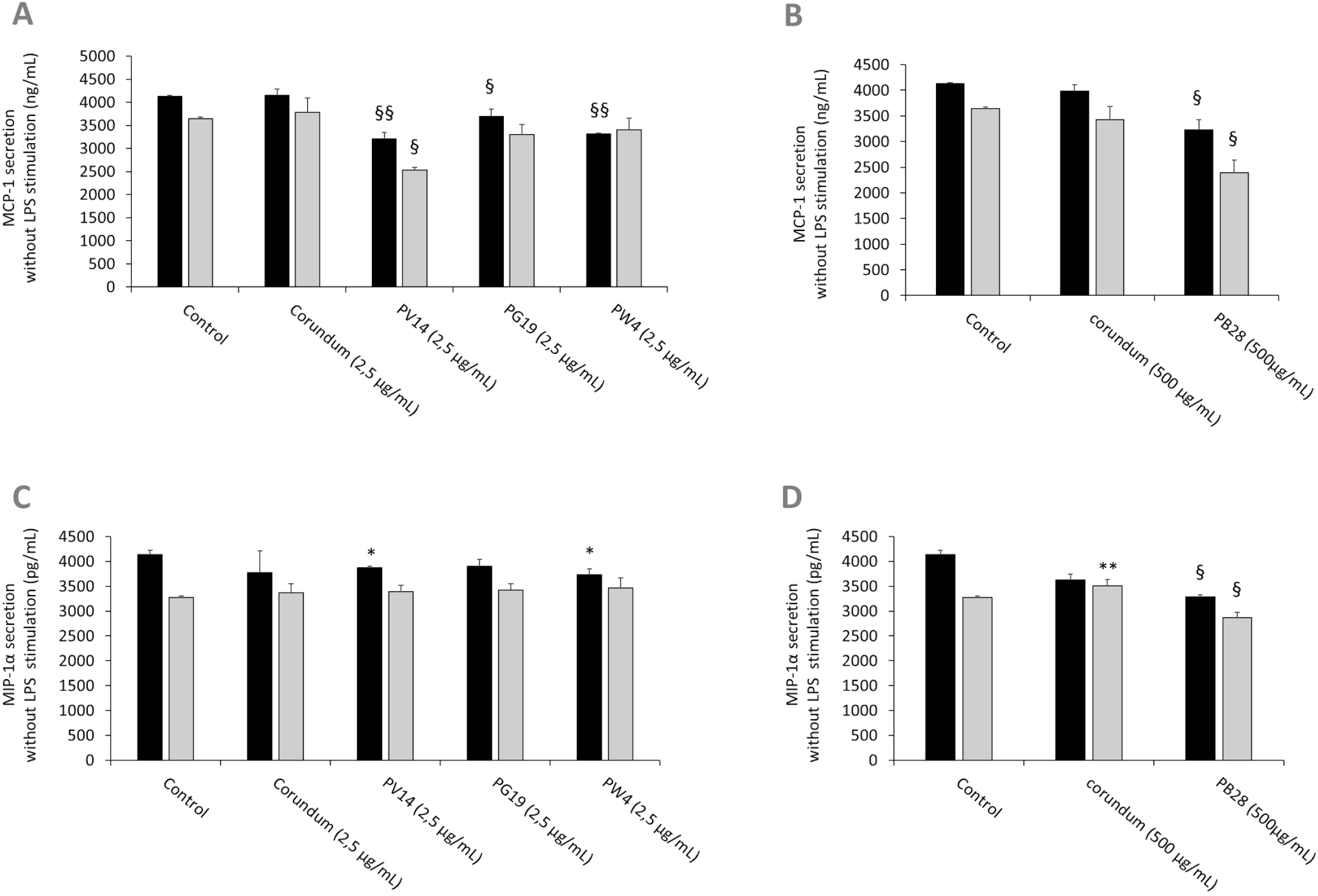
chemoattractant cytokines secretion without LPS stimulation. **Black bars: acute exposure (24 hours). Grey bars: exposure for 24 hours followed by a 3-day post-exposure recovery period**. Panel A: MCP-1 secretion. Conditions with 2.5 µg/ml of particle suspension. Panel B: MCP-1 secretion. Conditions with 500 µg/ml of particle suspension. Panel C: MIP-1α secretion. Conditions with 2.5 µg/ml of particle suspension. Panel D: MIP-1α secretion. Condition with 500 µg/ml of particle suspension. Significance of symbols: *: different from negative control only; §: different from both negative and corundum controls. Number of symbols: 1: p<0.05; 2: p<0.01.

In the presence of LPS (Figure 13), macrophages responded to this stimulation by increasing strongly the secretion of MCP-1 and MIP-1α. However, during acute exposure, we observed for PW4 and PB28 a significant decrease in MCP-1 secretion of 44656 pg/mL and 38404 pg/mL respectively compared to their respective controls (Control without treatment = 73587 pg/mL, Corundum (2.5 µg/mL) = 67899 pg/mL and Corundum (500 µg/mL) = 73433 pg/mL). For PW4, the effect is temporary and did not persist during recovery exposures, contrary to PB28 where the decrease remained strong (9782 pg/mL) compared to its respective controls (37652 pg/mL for control without treatment and for 28111 pg/mL for corundum (500 µg/mL)). In this experiment, we also noticed that PV14 did not seem to have an immediate effect on MCP-1 secretion but tended to display a delayed effect. The decrease is only visible in recovery exposure but this tendency is only significant compared to control without treatment. Concerning the secretion of MIP-1α in the presence of LPS, only PB28 induced an effect in recovery exposure. Indeed, a significant decrease of approximately 100000 pg/mL compared to its controls was observed. thus, PB28 decreased MIP-1α secretion in a delayed manner.

**Figure 13.**
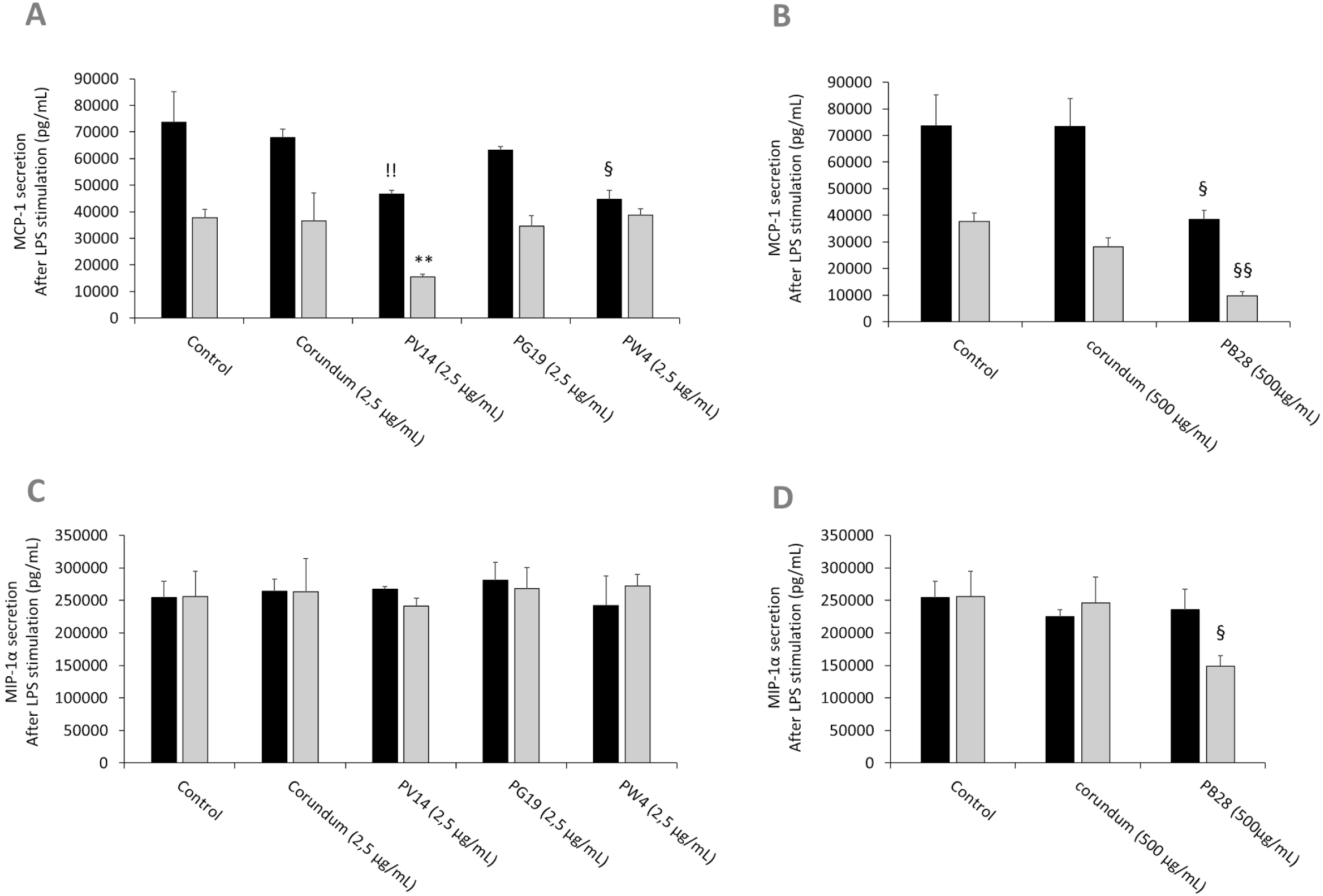
chemoattractant cytokines secretion after LPS stimulation. **Black bars: acute exposure (24 hours). Grey bars: exposure for 24 hours followed by a 3-day post-exposure recovery period**. Panel A: MCP-1 secretion. Conditions with 2.5 µg/ml of particle suspension. Panel B: MCP-1 secretion. Conditions with 500 µg/ml of particle suspension. Panel C: MIP-1α secretion. Conditions with 2.5 µg/ml of particle suspension. Panel D: MIP-1α secretion. Condition with 500 µg/ml of particle suspension. Significance of symbols: *: different from negative control only; §: different from both negative and corundum controls; !: different from corundum control only. Number of symbols: 1: p<0.05; 2: p<0.01.

### 3.7. Pigments increase the proportion of CD11b-negative macrophages

In order to know if pigments induced changes to the profiles of macrophages, we screened CD11b (Figures 14A and 14B), an integrin classically associated with macrophages and dendritic cells, which contributes to cell adhesion and immunological tolerance. We thus investigated the proportion of cells becoming negative for the presence of CD11b at their surface. The results in Figure 14A showed that, under acute exposure, there was no change in the presence of CD11b at the surface of cells in the presence of PV14 (2.5 µg/ml), PG19 (2.5 µg/ml), and PW4 (2.5 µg/mL) in comparison to control without treatment. A significant increase in CD11b-negative cells (19%) was observed for PG19 (2.5 µg/mL) in comparison to corundum control (2.5 µg/mL) (15%). However, we observed a slightly significant increase in CD11b-negative cells with a higher concentration of corundum (500 µg/ml) (16%) in comparison to control without treatment (11%) (Figure 14B). Moreover, the percentage of CD11b-negative cells increased dramatically in the case of PB28 (500 µg/ml) to reach 62% (Figure 14B).

**Figure 14.**
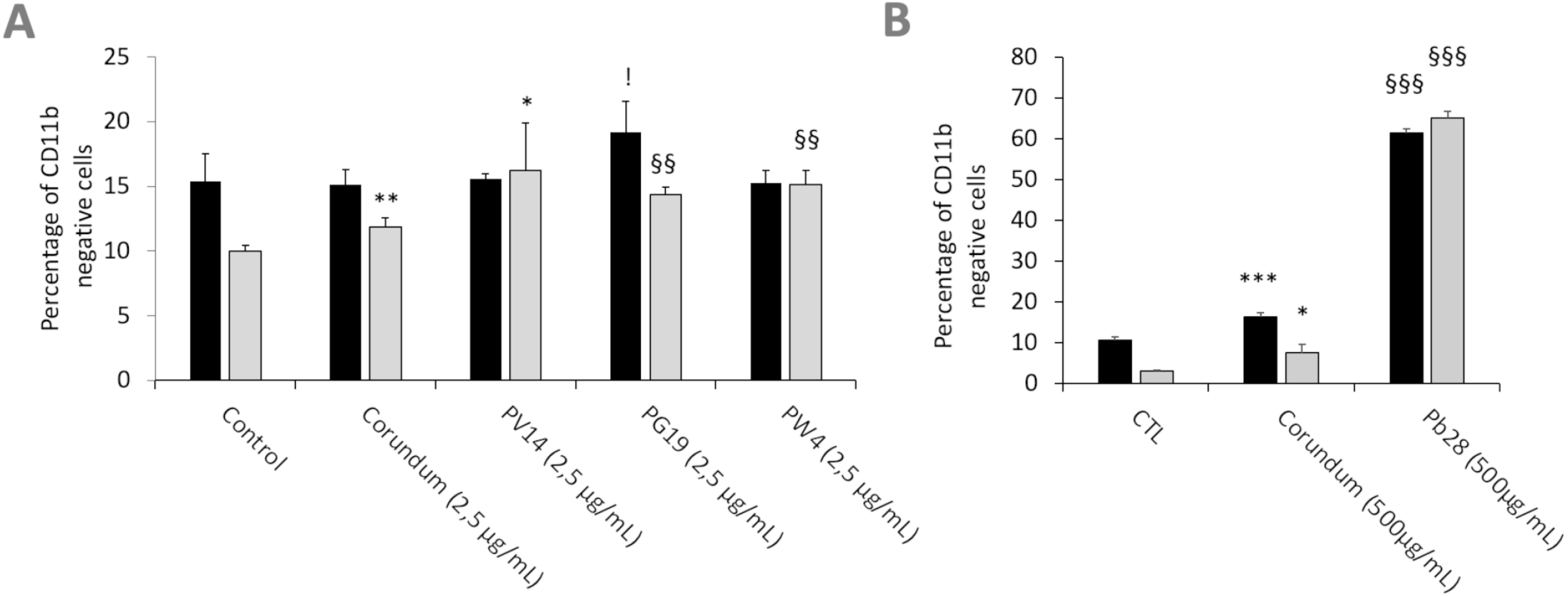
CD11b cell surface marking. **Black bars: acute exposure (24 hours). Grey bars: exposure for 24 hours followed by a3-day post-exposure recovery period**. Panel A: percentage of CD11b-cells in comparison to unlabeled control cells. Conditions with 2.5 µg/ml of particle suspension. Panel B: percentage of CD11b-cells in comparison to unlabeled control cells. Conditions with 500 µg/ml of particle suspension. Conditions with 500 µg/ml of particle suspension. Significance of symbols: *: different from negative control only; §: different from both negative and corundum controls; !: different from corundum control only. Number of symbols: 1: p<0.05; 2: p<0.01; 3: p<0.001.

This increase in the proportion of CD11b-negative cells was observed at the end of the recovery period, for both the high concentration of corundum and PB28 (65% of CD11b-negative cells in comparison to controls without treatment (3%) and corundum (500 µg/mL) (8%). This suggests that PB28 has a highly sustained effect (Figure 14B). Moreover, the increase became significant after recovery exposure in the case of the cells exposed to PG19 or PW4 (14 and 15% of CD11b negative cells instead of 10 or 12% for controls). PV14 (2.5 µg/mL) could have a small effect compared to the negative control, but not to the matched corundum control (Figure 14A). Thus, even if the effect was smaller than in the case of PB28 (500 µg/mL), the results suggested that PG19 (2.5 µg/mL) and PW4 (2.5 µg/mL) had a slight but significant delayed effect.

## 4. Discussion

In any nanotoxicology context, the first question to be addressed is the relevance of the dose to be tested. For tattoos, the injected dose of pigments is reported to be in between 0.5 and 0.9 mg/cm2, with an estimated mean of 2.5 mg/cm2 [17]. If it is assumed that all of these pigments are collected by dermal macrophages [21, 22], then this surface dose can be transposed to the pure macrophage culture. When transposed to a 12-well plate with 1 ml of culture medium per well and a culture area of 3.8 cm2, this amount to a dose close to 9.5 mg/ml of pigments. Metal analyses of tattoo inks have suggested that they often contain a mixture of pigments [40], so that the total pigment dose cannot be considered to be composed of only one pigment. But even considering this dilution phenomenon, the doses used in the present study cannot be considered to be exaggerated.

Because tattoo inks are poorly regulated, their composition under the same trade name may evolve over time. In this respect, the comparison of the metals contained in tattoo inks between 2009 and 2020 [40, 69] shows a reduction in toxic metals such as cobalt for example. Despite this positive trend, it should not be forgotten that data may be lacking regarding the people who were tattooed in the early 2000s, and therefore, regarding the potential long-term effects of these metal-containing pigments. Thus, the relevance of examining the effects of cobalt pigments on macrophages should not be underestimated. Even though they are often more expensive than organic pigments, mineral pigments show a high resistance to light, and degrade only by releasing their constituent metal ions, contrary to organic pigments that can have complex degradation pathways (e.g., in [70, 71]).

With the above considerations in mind, we decided to investigate the responses of macrophages to three different cobalt-based pigments, and one zinc-based pigment, which served as a comparison for PG19. Indeed, the results of ICP-AES showed the presence of a high quantity of zinc besides cobalt for PG19. Their toxicity is clearly split between toxic pigments (Pigment Violet 14, Pigment Green 19, and Pigment White 4), and low-toxicity pigments (Pigment Blue 28) leading to a very high LD20 concentration. As the latter could be used at high concentrations (i.e., close to 1 mg/ml), the possibility of macrophage being overloaded [72, 73] could not be ruled out. In any case, compared to the untreated negative control, the onset of phagocytosis due to the presence of particulate material is likely to induce changes to macrophage physiology, which should not be mistaken for specific effects induced by the cobalt pigments. We thus used the negative corundum control [74], at both low and high concentrations, in order to take these potential effects into account.

Furthermore, as the adverse effects of cobalt and/or zinc can be expected to occur via a progressive release in ionic form, we decided to take this parameter into account by using an exposure-recovery scheme, in which the degradation of the pigments inside the macrophage is likely to start to occur even for poorly-degradable pigments. For more degradable pigments, this degradation may occur even during the exposure period itself.

Indeed, one of the reasons for the low toxicity of pigments resides in their very low solubility in water, and thus, in biological media, so that no release of toxic components should occur. Mineral pigments known to be toxic, such as lead white (basic lead carbonate) and vermillion (mercuric sulfide) dissolve slowly in biological media, leading to a slow release of toxic heavy metal ions. In fact, biological media are more complex than plain water. Assuming that pigment particles end up, at least in part, in macrophages’ phagolysosomes, these particles will encounter a mildly acidic medium (pH 5) containing a high concentration of proteins (comprising enzymes) that bind many metal ions with high affinity. This may explain why water-insoluble pigments may prove soluble to some extent in biological media. This description of the biological media also suggests that pigment biosolubility will depend on the structure of each pigment. The results on cobalt pigments show a clear correlation between the results obtained in strongly acidic mineralization and the results obtained in vivo. Pigments and minerals that are poorly soluble in acidic media, such as corundum or PB28, are also less easily dissolved in cells (0.01% after 24 hours) while pigments that dissolve more easily in acidic media, such as PV14, also dissolve more easily in cells (ca. 2% dissolved in cells after 24 hours). This is reflected in the toxicity of the pigments, as more soluble pigments show a higher toxicity than poorly soluble pigments (LD20 of 2.5 µg/ml for PV14 compared to 500 µg/ml for PB28). The data are more difficult to interpret for PG19, as it contains too little cobalt to distinguish it from the background levels.

Because of this difference in solubility, the biological effects of the pigments are likely to be a mixture of purely nano- or microparticulate effects along with effects linked to ion release. This is why we systematically included corundum as a particulate control. Indeed, it has been shown to exhibit minimal toxicity and no pro-inflammatory effects, which makes it of great interest to be studied in macrophage biology [74].

One of the effects that can be expected to be dependent on particle concentration is phagocytosis, which is linked to the well-known overload effect [72, 73]. However, our results showed that phagocytosis was also impacted by a low quantity of internalized particles. This slight decrease is probably not due to an overload effect but can be explained by the release of soluble cobalt or zinc ions via the lysosomal or phagolysosomal process. At a higher concentration, we observed a correlation between the overload effect and the effects detected in the present study. Interestingly enough, the highly-corundum loaded cells recovered a normal phagocytic activity after a few days while the cells exposed to an equal concentration of a cobalt-containing pigment with the same crystalline structure (spinel) did not, which clearly shows that cobalt has a delayed effect. This persisting effect was also observed for a lower concentration of pigment, but only for those including zinc, such as PG19 and PW4, suggesting a specific effect of zinc. All particles also induced a weak pro-inflammatory effect. This is also true in the case of corundum (at high concentration), while this effect had not been detected previously in an indirect assay, but in serum-free culture conditions [74]. These results were first observed with a NO assay and already revealed an overall persistent pro-inflammatory effect of PV14, PG19 and PW4. Thus, the engulfment of cobalt and/or zinc (nano)particles present in some pigments could induce a slight, long-term inflammation. No synergistic effect was identified in the case of a mixture of zinc and cobalt. Indeed, the intrinsic long-term pro-inflammatory effect was similar between PV14, PG19, and PW4. On the contrary, the mixture of cobalt and aluminum did not induce persisting effect at high quantity while corundum (which is composed of aluminum oxide) induced a persisting effect at high concentration. For TNF, this effect was essentially transient. However, a decrease was noted after recovery exposure in the case of the cells exposed to PB28 at high concentration. This result is in line with those observed via the NO assay and the reason is probably linked to the presence of soluble cobalt + aluminum ions release in cells.

This scheme was not observed for interleukin 6: no significant sustained stimulation was observed except for the higher concentration of particles (500 µg/mL for corundum and PB28). Like phagocytosis, the intrinsic transient and persisting effects of (nano)particles on IL-6 are due to an overload of internalized materials, but it can also be aggravated by the long-term release of cobalt ions after a high quantity of PB28 has been engulfed.

To go further into the question of the intrinsic effects of cobalt-containing pigments on macrophages, we also investigated their genotoxicity. Interestingly, the highly soluble and toxic pigments did not show a detectable genotoxicity, even though the amount of intracellular soluble cobalt is well above the concentration used as soluble cobalt on total leukocytes (35-65 fg Co/cell in this work, based on a protein content of 300 pg protein/cell [75], compared to the 5-8 fg Co/cell used on leukocytes) [44]. This result is in line with the established resistance of macrophages to genotoxicity, as opposed to lymphocytes, which account for the vast majority of mononuclear leukocytes, and are highly sensitive to genotoxic stresses. Interestingly, a genotoxic effect was observed with high concentrations of particles, whether in the case of corundum or PB28. This effect was transient for corundum but sustained for PB28. This effect may be correlated with either the release of soluble cobalt and aluminum (Table 2) or an intrinsic effect of some insoluble particles (in this particular case, an alloy of aluminum and cobalt), as described in the case of titanium dioxide (reviewed in [76]). Indeed, an overload of particles in cells could disturb antioxidative and DNA repair processes [77], or may alter directly the cell division process, as suggested by our data (S4)

In addition to these effects, we investigated other effects of cobalt-containing pigments on the functionality of macrophages. We first investigated the response to a LPS challenge which would mimic an antibacterial response. We did not observe any significant change to the secretion of IL-6. Regarding the secretion of TNF, we only observed a moderate increase for PG19 and only immediately after exposure. The most striking fact is the general decrease in the responses of TNF and IL-6 after the recovery period. Coupled with the effects on phagocytosis, these results suggest that macrophages become less efficient in fighting a bacterial infection when loaded with cobaltand/or zinc-containing pigments. Corundum also induced a decrease of TNF and IL-6 (though lower than PB28) and only at a high concentration (500 µg/mL). Thus, for PB28, an overload effect is probably combined with the dissolution of the alloy of cobalt and aluminum. This sort of effect was not observed in the case of macrophages treated with amorphous silica [54]. It was observed, however, in the case of macrophages treated with silver nanoparticles [53], which also release toxic metallic ions during the recovery phase. When compared to results obtained under chronic exposure to silver nanoparticles [56], the results of this study on cobalt and/or zinc-containing pigments suggest that this decrease in performance may last over time, a fact which is to be taken into account in the perspective of the long-term effects of tattoos, where a higher susceptibility to infections (which may be the cause of certain chronic skin diseases) has been described [4,30,31].

Concerning chemokines (or chemoattractant cytokines) such as MCP-1 and MIP-1α involved in the recruitment of leukocytes. it has been shown that they may be involved in multiple inflammatory pathologies [79, 80] and have several functions other than chemokines, which are not yet all elucidated. For example, it is now demonstrated that MCP-1 also plays a role in cell adhesion by promoting the presence of integrins such as CD11b [81]. MCP-1 increases the phagocytic activity of macrophages and influences macrophage polarization and cell survival [79, 82]. It was therefore logical to focus on the perturbations that the selected inks can induce on this chemokine. PV14 (2,5 µg/mL) and PB28 (2,5 µg/mL) induce a persistent decrease in MCP-1 secretion by macrophages both in the absence and presence of LPS. This suggests a reduced ability to recruit monocytes to the skin site with these inks during infections. To a lesser extent, MCP-1 secretion is also decreased in the absence of bacterial stimuli, which also suggests that the removal of cell debris and apoptotic bodies or even foreign bodies may be less efficient during renewal (the skin is renewed approximately every 20 to 30 days depending on age [83] and tissue repair stages. Moreover, it is logical to wonder about the renewal capacity of skin resident macrophages if the recruitment of circulating monocytes is of lesser quality. A greater susceptibility to infections as well as a less efficient skin renewal may possibly explain a greater susceptibility to chronic skin inflammations (especially if the conditions are favorable for the development of skin diseases). Concerning MIP-1α, this chemokine is normally strongly secreted in the presence of LPS. This is observed in the majority of the results obtained with the pigments except for PB28 (500 µg/mL) which seems to lower MIP 1α levels in a prolonged way. This was also observed in the absence of LPS stimulation. This reinforces the interpretation of the results obtained with MCP-1. In the presence of PB28, macrophages appear to be less able to enroll other immune cells (such as macrophages or CD8 T lymphocytes) at the infected or inflamed site to stop pathogens or clean up the damaged skin (depending on the stage of inflammation). Thus, these results prove a more or less important disorder in the production of cytokines and chemokines depending on the pigment tested. If different stages of inflammation are deregulated, it is therefore logical to imagine that there could be a greater susceptibility to skin diseases in the long term. Moreover, Similar to MCP-1, MIP-1α plays a role in the regulation of CD11b integrin on macrophages surface [81].

Finally, we also tested CD11b, a cell-surface marker important for the good functioning of macrophages. Indeed, CD11b is also part of the complement receptor 3, and, as such, plays a role in phagocytosis for complement-opsonized pathogens [84]. Thus, CD11b-negative cells are likely to be impaired, and thus, less efficient for the elimination of pathogens.

Last but not least, CD11b also plays a role in immune regulation as a negative regulator of immune responses [85, 86]. Thus, the absence of CD11b results in a higher T-cell activation [87] and may result in a reduction of tolerance and deregulated inflammatory response [88]. Moreover, the activation of CD11b, which is linked to the anti-tumor immune response [89], will be impossible in CD11b-negative cells, and may contribute to a lower anti-tumoral activity. Thus, the increase in the CD11b-negative population that we observed on pigment-treated cells, especially after the recovery period, may be linked to the observed inflammatory complications linked to tattoos [4,6,7]

Overall, the macrophages treated with zinc and cobalt-containing pigments appear less efficient against pathogens, may be less efficient against tumors and they also seem to have a more proinflammatory basal phenotype. Moreover, based on results, macrophages should be less efficient in their key role in tissue renewal (elimination of cellular debris and apoptotic cells via efferocytosis). Indeed, a large fraction of monocytes can potentially be cleared as a by-product of immune surveillance. Of course, the effects are likely to be moderate, and it can be argued that the effects that we detected over a few days may normalize over time. However, our results on the persistence of these effects, coupled with the now-established capture-death-recapture macrophage cycle following the injection of tattoo pigments [22, 23], point towards a self-sustaining process which alters macrophages on the long term.

Although mineral pigments (such as cobalt or zinc nano-and microparticles) are now seldom used in tattoo inks, the fact that people may have received such pigments in the past [40] as well as the possibility that other pigments may also alter the functions of macrophages, possibly via completely different mechanisms, point to the need for more research in the field of sustained/delayed effects of pigments.

Moreover, even if the results are not exactly the same as those obtained in this paper, other articles confirm that the presence of cobalt ions (and also of other metals such as copper) induces changes to the functionalities of macrophages, in particular their pro-or antiinflammatory profile. The differences observed between these works and ours are probably related to the different form of the cobalt elements used, since in our work, the metal elements were initially present as particles and not directly as ions, as in [90].

## 5. Conclusion

In conclusion (Figure 15), the pigments tested contain nano-or sub-micrometer objects, and not only microparticles. Overall, looking at the data regarding the secretion of NO, IL-6 and TNF, the pigments tested seem to have a moderate intrinsic pro-inflammatory effect. This effect, however, may persist over time. Concerning the body’s defense mechanisms, the macrophages which have internalized the cobalt- or zinc-based pigments have a lower ability to respond to a bacterial infection (e.g., skin infection). Indeed, the phagocytic capacity of macrophages and their capacity to respond to inflammatory stimuli decrease, and this effect is persistent. This idea is reinforced by the additional presence of CD11b-negative cells. Moreover, and mostly observed for PB28 pigment, chemokines MCP-1 and MIP-1α are decreased persistently in absence or presence of bacterial stimuli. Following this study, the question arises as to whether the inks containing cobalt and/or zinc micro- and nano-objects are safe or not. It is legitimate to wonder if they can generate localized or generalized alterations of immunoregulation and weaken the body’s defense mechanisms against bacterial pathogens and their ability to recognize cancer cells. A suspicion of disturbances during tissue regeneration can also be considered. This can be explained by a decrease in the phagocytic capacity (and thus in efferocytosis) of the macrophages present at the tattoo site as well as a local reduction in mediators allowing the recruitment of other cells of the immune system, including the macrophages themselves. Consequently, the cellular homeostasis and immunological surveillance which is necessarily arbitrated by the cells of the innate then adaptive immunity is necessarily disturbed. The obtained results can thus give first explanations concerning the disturbances of certain molecular and cellular mechanisms associated with the emergence of chronic cutaneous diseases in tattooed patients [4, 6]. The effects which were observed seem to have been reinforced by overload effects (for high concentrations of pigments), and are proportionally linked to the release of free ions during the dissolution in the phagolysosomes.

**Figure 15.**
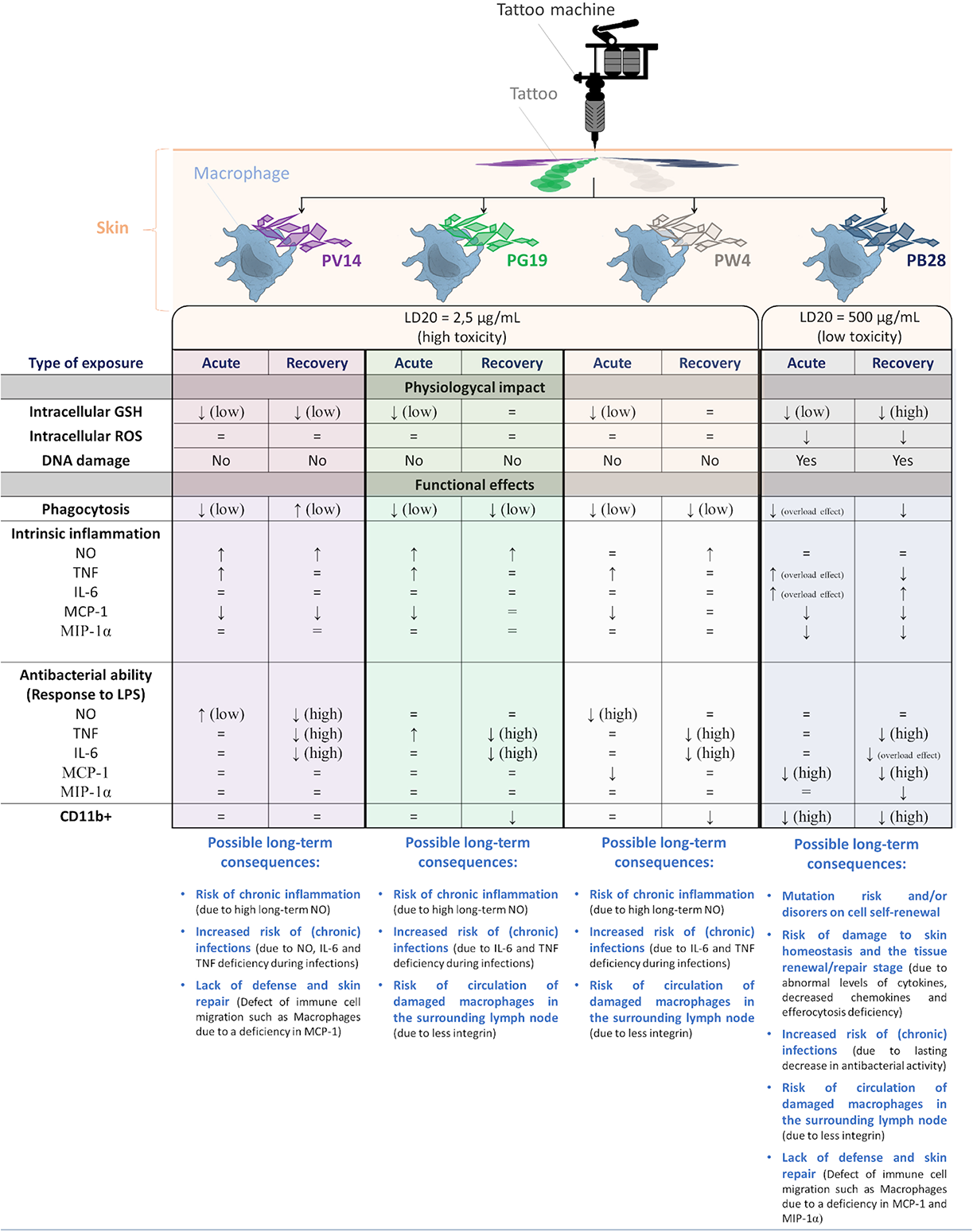
– Summary figure of the results obtained and possible consequences on the skin for the different inks tested. Significance of symbols:= no differences ; ↓ decrease in comparison to controls ; ↑ increase in comparison to controls.

Finally, a better understanding of the disturbances of the inflammatory mechanisms (cellular and molecular disorders) in the presence of tattoos will allow a better treatment of the patients having contracted chronic pathologies of the skin (maybe generalized) following the tattoos. The treatments are still not well known.

## Supporting information

supdata for manuscript

## Supplementary data

**Supplementary data 1.**
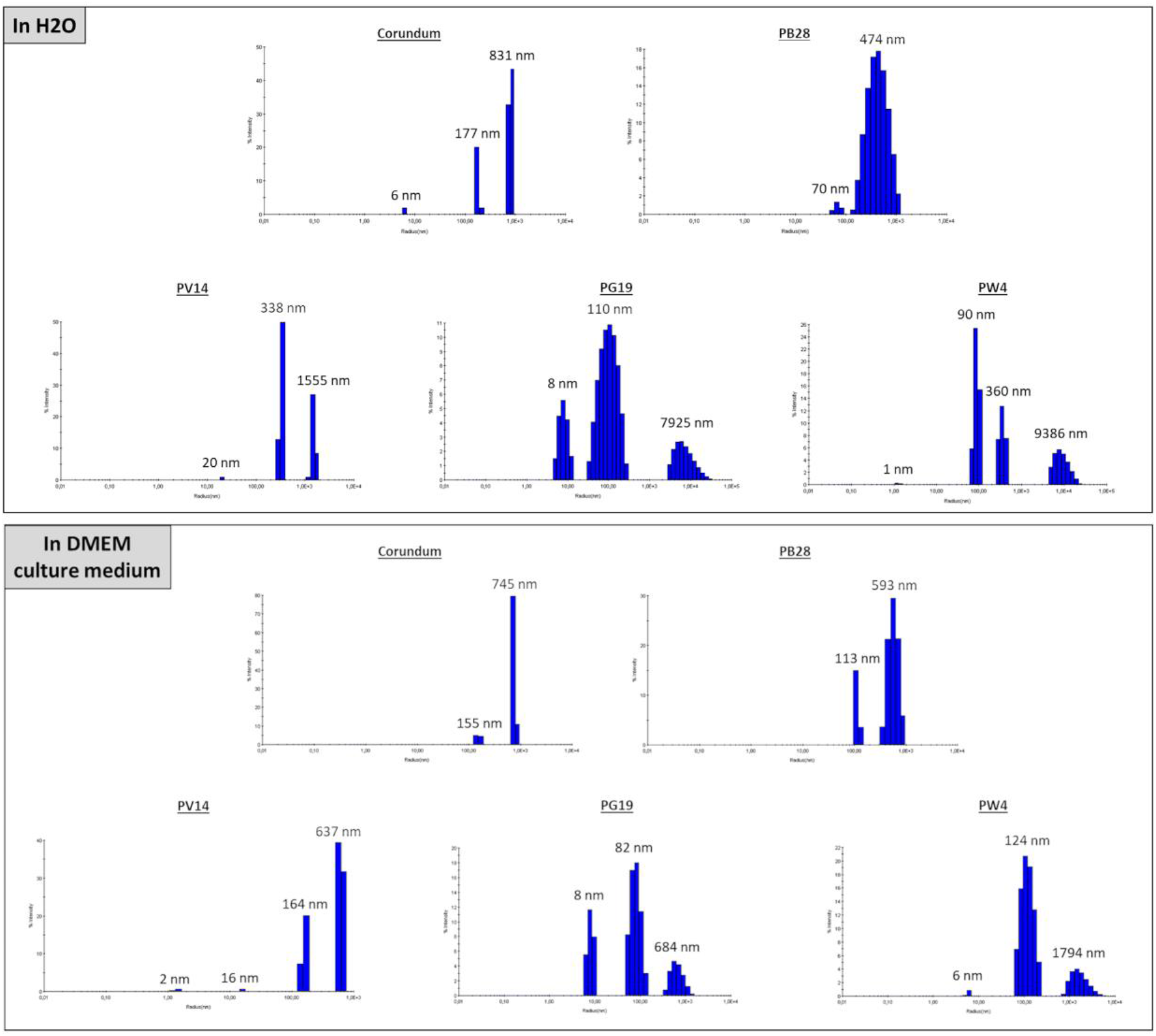
Example of hydrodynamic diameter measurement (screen print of raw data) by DLS in H_2_O and DMEM culture medium.

**Supplementary data 2.**
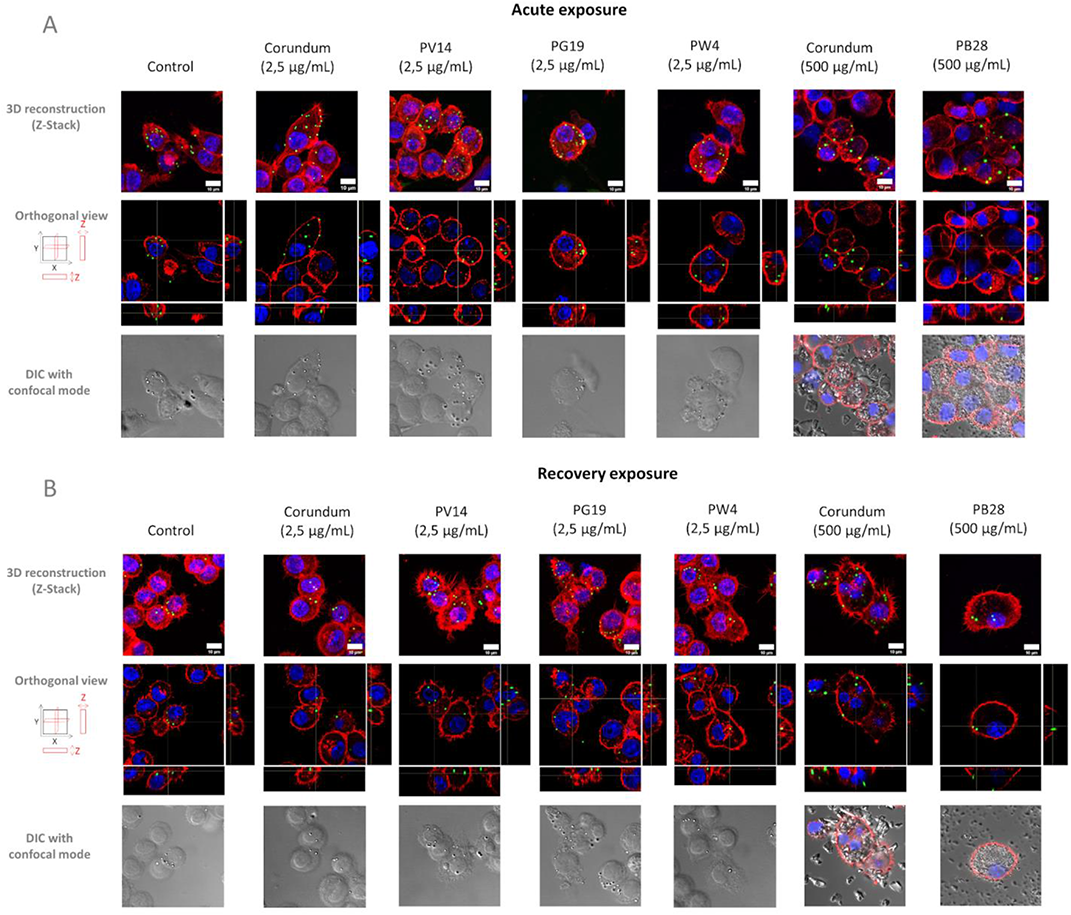
Visualization of phagocytic activity by confocal microscopy. Pannel A- Acute exposure of macrophages to pigments. Pannel B – End of recovery exposure of macrophages to pigment. Red color = Actin filament visualized with phalloidin-Atto 560. Blue color = Nucleus visualized with Dapi. Green color = Fluorescent yellow/green carboxylate-modified-polystyren beads (1 µm diameter). Grey color = differential interference contrast (DIC). Scale bar = 10 µm.

**Supplementary data 3.**
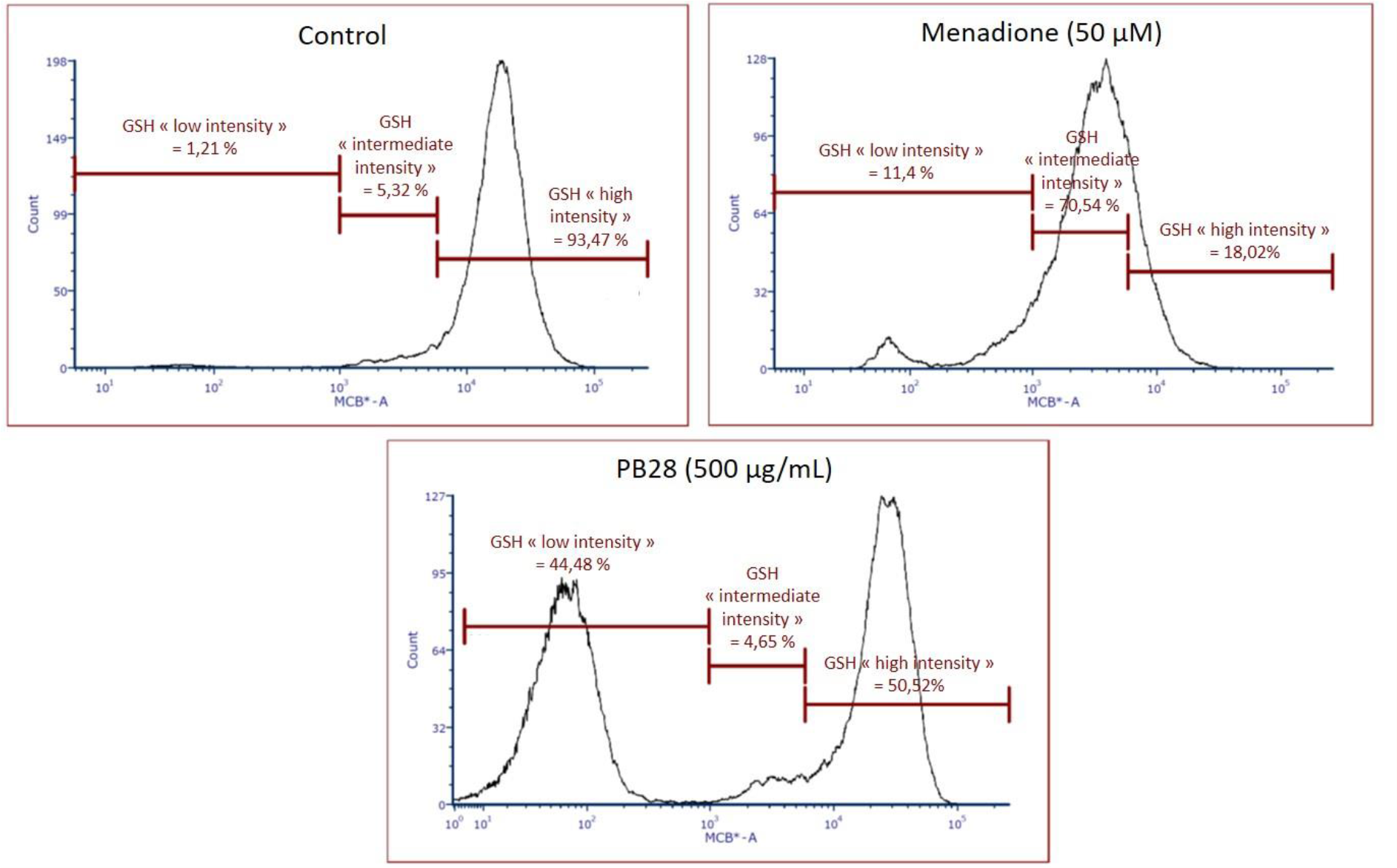
Examples of GSH results obtained via flow cytometry (Raw results of recovery exposure).

**Supplementary data 4.**
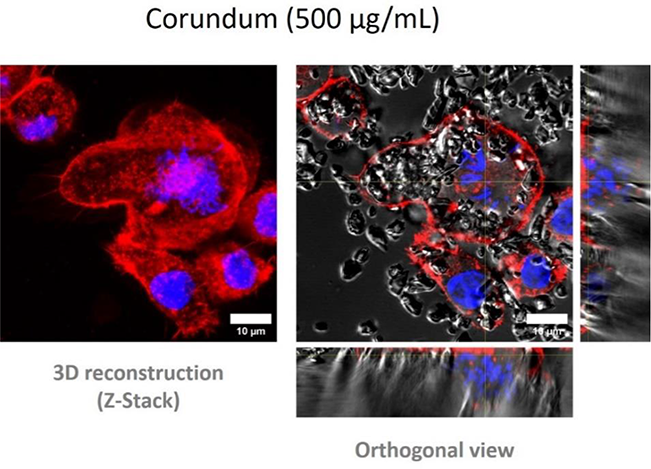
Confocal microscopy. Observation of actin conformation and nuclear integrity. Red = Actin (in red) stained with phalloidin-Atto 560. DNA (in blue) stained with Dapi. Grey = Corundum particles observed in confocal DIC. Scale bar = 10 µm.

## Author contributions

JD and BD performed experiments on the functional effects of pigments on macrophages and ICP-AES measurements. MD and MC performed the genotoxicity assays. JP trained the experimenters at the ICP-AES. TR analyzed the ICP-AES results. DF performed TEM microscopy visualization. VCF acquired and analyzed flow cytometry experiments. BD and TR supervised the overall research. The manuscript was written by TR, BD, and to a lesser extent, JD. All co-authors approved the final version of the manuscript.

## References

1. Deter-Wolf, A. The Material Culture and Middle Stone Age Origins of Ancient Tattooing. In Tattoos and Body Modifications in Antiquity. Proceedings of the sessions at the EAA annual meetings in The Hague and Oslo, 2010/11; Zurich Studies in Archaeology; 2013; pp. 15–26.

2. Dorfer, L.; Moser, M.; Bahr, F.; Spindler, K.; Egarter-Vigl, E.; Giullén, S.; Dohr, G.; Kenner, T. A Medical Report from the Stone Age? Lancet 1999, 354, 1023–1025, doi:10.1016/S0140-6736(98)12242-0.

3. Klügl, I.; Hiller, K.-A.; Landthaler, M.; Bäumler, W. Incidence of Health Problems Associated with Tattooed Skin: A Nation-Wide Survey in German-Speaking Countries. Dermatology 2010, 221, 43–50, doi:10.1159/000292627.

4. Islam, P.S.; Chang, C.; Selmi, C.; Generali, E.; Huntley, A.; Teuber, S.S.; Gershwin, M.E. Medical Complications of Tattoos: A Comprehensive Review. Clinic Rev Allerg Immunol 2016, 50, 273–286, doi:10.1007/s12016-016-8532-0.

5. Laumann, A.E.; Derick, A.J. Tattoos and Body Piercings in the United States: A National Data Set. Journal of the American Academy of Dermatology 2006, 55, 413–421, doi:10.1016/j.jaad.2006.03.026.

6. van der Bent, S.A.S.; Rauwerdink, D.; Oyen, E.M.M.; Maijer, K.I.; Rustemeyer, T.; Wolkerstorfer, A. Complications of Tattoos and Permanent Makeup: Overview and Analysis of 308 Cases. J of Cosmetic Dermatology 2021, 20, 3630–3641, doi:10.1111/jocd.14498.

7. Yoong, C.; Vun, Y.Y.; Spelman, L.; Muir, J. True Blue Football Fan: Tattoo Reaction Confined to Blue Pigment. Australasian Journal of Dermatology 2010, 51, 21–22, doi:10.1111/j.1440-0960.2009.00608.x.

8. Lerche, C.M.; Heerfordt, I.M.; Serup, J.; Poulsen, T.; Wulf, H.C. Red Tattoos, Ultraviolet Radiation and Skin Cancer in Mice. Exp Dermatol 2017, 26, 1091–1096, doi:10.1111/exd.13383.

9. Pabst, M.A.; Letofsky-Papst, I.; Moser, M.; Spindler, K.; Bock, E.; Wilhelm, P.; Leopold Dorfer, M.D.; Geigl, J.B.; Auer, M.; Speicher, M.R.;, et al. Different Staining Substances Were Used in Decorative and Therapeutic Tattoos in a 1000-Year-Old Peruvian Mummy. Journal of Archaeological Science 2010, 37, 3256–3262, doi:10.1016/j.jas.2010.07.026.

10. Laux, P.; Tralau, T.; Tentschert, J.; Blume, A.; Dahouk, S.A.; Bäumler, W.; Bernstein, E.; Bocca, B.; Alimonti, A.; Colebrook, H.;, et al. A Medical-Toxicological View of Tattooing. The Lancet 2016, 387, 395–402, doi:10.1016/S0140-6736(15)60215-X.

11. COMMISSION REGULATION (EU) 2020/2081 Amending Annex XVII to Regulation (EC) No 1907/2006 of the European Parliament and of the Council Concerning the Registration, Evaluation, Authorisation and Restriction of Chemicals (REACH) as Regards Substances in Tattoo Inks or Permanent Make-Up 2020.

12. Maarouf, M.; Saberian, C.; Segal, R.J.; Shi, V.Y. A New Era For Tattoos, with New Potential Complications. J Clin Aesthet Dermatol 2019, 12, 37–38.

13. Maarouf, M.; Saberian, C.; Segal, R.J.; Shi, V.Y. A New Era For Tattoos, with New Potential Complications. J Clin Aesthet Dermatol 2019, 12, 37–38.

14. Paprottka, F.J.; Krezdorn, N.; Narwan, M.; Turk, M.; Sorg, H.; Noah, E.M.; Hebebrand, D. Trendy Tattoos-Maybe a Serious Health Risk? Aesthetic Plast Surg 2018, 42, 310– 321, doi:10.1007/s00266-017-1002-0.

15. Jacob, C.I. Tattoo-Associated Dermatoses: A Case Report and Review of the Literature. Dermatol Surg 2002, 28, 962–965, doi:10.1046/j.1524-4725.2002.02066.x.

16. Forbat, E.; Al-Niaimi, F. Patterns of Reactions to Red Pigment Tattoo and Treatment Methods. Dermatol Ther (Heidelb) 2016, 6, 13–23, doi:10.1007/s13555-016-0104-y.

17. Baumler, W. Absorption, Distribution, Metabolism and Excretion of Tattoo Colorants and Ingredients in Mouse and Man: The Known and the Unknown. In Current Problems in Dermatology; Serup, J., Kluger, N., Baumler, W., Eds.; S. Karger AG, 2015; Vol. 48, pp. 176–184 ISBN 978-3-318-02776-1.

18. Regensburger, J.; Lehner, K.; Maisch, T.; Vasold, R.; Santarelli, F.; Engel, E.; Gollmer, A.; König, B.; Landthaler, M.; Bäumler, W. Tattoo Inks Contain Polycyclic Aromatic Hydrocarbons That Additionally Generate Deleterious Singlet Oxygen. Exp Dermatol 2010, 19, e275–281, doi:10.1111/j.1600-0625.2010.01068.x.

19. Kluger, N. Cutaneous Complications Related to Permanent Decorative Tattooing. Expert Rev Clin Immunol 2010, 6, 363–371, doi:10.1586/eci.10.10.

20. Wenzel, S.M.; Rittmann, I.; Landthaler, M.; Bäumler, W. Adverse Reactions after Tattooing: Review of the Literature and Comparison to Results of a Survey. Dermatology 2013, 226, 138–147, doi:10.1159/000346943.

21. Kluegl, I.; Hiller, K.-A.; Landthaler, M.; Baeumler, W. Incidence of Health Problems Associated with Tattooed Skin: A Nation-Wide Survey in German-Speaking Countries. Dermatology 2010, 221, 43–50, doi:10.1159/000292627.

22. Baranska, A.; Shawket, A.; Jouve, M.; Baratin, M.; Malosse, C.; Voluzan, O.; Vu Manh, T.-P.; Fiore, F.; Bajénoff, M.; Benaroch, P.;, et al. Unveiling Skin Macrophage Dynamics Explains Both Tattoo Persistence and Strenuous Removal. J Exp Med 2018, 215, 1115– 1133, doi:10.1084/jem.20171608.

23. Collin, M. Death, Eaters, and Dark Marks. J Exp Med 2018, 215, 1005–1006, doi:10.1084/jem.20180311.

24. Navegantes, K.C.; de Souza Gomes, R.; Pereira, P.A.T.; Czaikoski, P.G.; Azevedo, C.H.M.; Monteiro, M.C. Immune Modulation of Some Autoimmune Diseases: The Critical Role of Macrophages and Neutrophils in the Innate and Adaptive Immunity. J Transl Med 2017, 15, 36, doi:10.1186/s12967-017-1141-8.

25. Chen, G.Y.; Nuñez, G. Sterile Inflammation: Sensing and Reacting to Damage. Nat Rev Immunol 2010, 10, 826–837, doi:10.1038/nri2873.

26. Chanmee, T.; Ontong, P.; Konno, K.; Itano, N. Tumor-Associated Macrophages as Major Players in the Tumor Microenvironment. Cancers (Basel) 2014, 6, 1670–1690, doi:10.3390/cancers6031670.

27. Kohout, J. MACROPHAGE REACTIVITY IN SKIN WINDOW OF SARCOIDOSIS PATIENTS. Ann NY Acad Sci 1976, 278, 201–203, doi:10.1111/j.1749-6632.1976.tb47030.x.

28. Isohisa, T.; Asai, J.; Kanemaru, M.; Arita, T.; Tsutsumi, M.; Kaneko, Y.; Arakawa, Y.; Wada, M.; Konishi, E.; Katoh, N. CD163-positive Macrophage Infiltration Predicts Systemic Involvement in Sarcoidosis. J Cutan Pathol 2020, 47, 584–591, doi:10.1111/cup.13675.

29. Standiford, T.J. Macrophage Polarization in Sarcoidosis: An Unexpected Accomplice? Am J Respir Cell Mol Biol 2019, 60, 9–10, doi:10.1165/rcmb.2018-0298ED.

30. Kasraie, S.; Werfel, T. Role of Macrophages in the Pathogenesis of Atopic Dermatitis. Mediators of Inflammation 2013, 2013, 1–15, doi:10.1155/2013/942375.

31. Forte, G.; Petrucci, F.; Cristaudo, A.; Bocca, B. Market Survey on Toxic Metals Contained in Tattoo Inks. Science of The Total Environment 2009, 407, 5997–6002, doi:10.1016/j.scitotenv.2009.08.034.

32. Chen, G.Y.; Nuñez, G. Sterile Inflammation: Sensing and Reacting to Damage. Nat Rev Immunol 2010, 10, 826–837, doi:10.1038/nri2873.

33. Navegantes, K.C.; de Souza Gomes, R.; Pereira, P.A.T.; Czaikoski, P.G.; Azevedo, C.H.M.; Monteiro, M.C. Immune Modulation of Some Autoimmune Diseases: The Critical Role of Macrophages and Neutrophils in the Innate and Adaptive Immunity. J Transl Med 2017, 15, 36, doi:10.1186/s12967-017-1141-8.

34. Schreiver, I.; Hesse, B.; Seim, C.; Castillo-Michel, H.; Villanova, J.; Laux, P.; Dreiack, N.; Penning, R.; Tucoulou, R.; Cotte, M.;, et al. Synchrotron-Based ν-XRF Mapping and μ-FTIR Microscopy Enable to Look into the Fate and Effects of Tattoo Pigments in Human Skin. Sci Rep 2017, 7, 11395, doi:10.1038/s41598-017-11721-z.

35. Nopajaroonsri, C.; Simon, G.T. Phagocytosis of Colloidal Carbon in a Lymph Node. Am J Pathol 1971, 65, 25–42.

36. Engel, E.; Vasold, R.; Santarelli, F.; Maisch, T.; Gopee, N.V.; Howard, P.C.; Landthaler, M.; Bäumler, W. Tattooing of Skin Results in Transportation and Light-Induced Decomposition of Tattoo Pigments--a First Quantification in Vivo Using a Mouse Model. Exp Dermatol 2010, 19, 54–60, doi:10.1111/j.1600-0625.2009.00925.x.

37. Lehner, K.; Santarelli, F.; Penning, R.; Vasold, R.; Engel, E.; Maisch, T.; Gastl, K.; König, B.; Landthaler, M.; Bäumler, W. The Decrease of Pigment Concentration in Red Tattooed Skin Years after Tattooing. J Eur Acad Dermatol Venereol 2011, 25, 1340–1345, doi:10.1111/j.1468-3083.2011.03987.x.

38. Khan, Z.; Combadière, C.; Authier, F.-J.; Itier, V.; Lux, F.; Exley, C.; Mahrouf-Yorgov, M.; Decrouy, X.; Moretto, P.; Tillement, O.;, et al. Slow CCL2-Dependent Translocation of Biopersistent Particles from Muscle to Brain. BMC Med 2013, 11, 99, doi:10.1186/1741-7015-11-99.

39. Gherardi, R.K.; Eidi, H.; Crépeaux, G.; Authier, F.J.; Cadusseau, J. Biopersistence and Brain Translocation of Aluminum Adjuvants of Vaccines. Front Neurol 2015, 6, 4, doi:10.3389/fneur.2015.00004.

40. Forte, G.; Petrucci, F.; Cristaudo, A.; Bocca, B. Market Survey on Toxic Metals Contained in Tattoo Inks. Science of The Total Environment 2009, 407, 5997–6002, doi:10.1016/j.scitotenv.2009.08.034.

41. Battistini, B.; Petrucci, F.; De Angelis, I.; Failla, C.M.; Bocca, B. Quantitative Analysis of Metals and Metal-Based Nano- and Submicron-Particles in Tattoo Inks. Chemosphere 2020, 245, 125667, doi:10.1016/j.chemosphere.2019.125667.

42. Simonsen, L.O.; Harbak, H.; Bennekou, P. Cobalt Metabolism and Toxicology--a Brief Update. Sci Total Environ 2012, 432, 210–215, doi:10.1016/j.scitotenv.2012.06.009.

43. Papageorgiou, I.; Brown, C.; Schins, R.; Singh, S.; Newson, R.; Davis, S.; Fisher, J.; Ingham, E.; Case, C.P. The Effect of Nano- and Micron-Sized Particles of Cobalt-Chromium Alloy on Human Fibroblasts in Vitro. Biomaterials 2007, 28, 2946–2958, doi:10.1016/j.biomaterials.2007.02.034.

44. Colognato, R.; Bonelli, A.; Ponti, J.; Farina, M.; Bergamaschi, E.; Sabbioni, E.; Migliore, L. Comparative Genotoxicity of Cobalt Nanoparticles and Ions on Human Peripheral Leukocytes in Vitro. Mutagenesis 2008, 23, 377–382, doi:10.1093/mutage/gen024.

45. Kwon, Y.-M.; Xia, Z.; Glyn-Jones, S.; Beard, D.; Gill, H.S.; Murray, D.W. Dose-Dependent Cytotoxicity of Clinically Relevant Cobalt Nanoparticles and Ions on Macrophages in Vitro. Biomed Mater 2009, 4, 025018, doi:10.1088/1748-6041/4/2/025018.

46. Kumanto, M.; Paukkeri, E.-L.; Nieminen, R.; Moilanen, E. Cobalt(II) Chloride Modifies the Phenotype of Macrophage Activation. Basic Clin Pharmacol Toxicol 2017, 121, 98– 105, doi:10.1111/bcpt.12773.

47. Schuster, B.E.; Roszell, L.E.; Murr, L.E.; Ramirez, D.A.; Demaree, J.D.; Klotz, B.R.; Rosencrance, A.B.; Dennis, W.E.; Bao, W.; Perkins, E.J.;, et al. In Vivo Corrosion, Tumor Outcome, and Microarray Gene Expression for Two Types of Muscle-Implanted Tungsten Alloys. Toxicology and Applied Pharmacology 2012, 265, 128–138, doi:10.1016/j.taap.2012.08.025.

48. Jarosz, M.; Olbert, M.; Wyszogrodzka, G.; Młyniec, K.; Librowski, T. Antioxidant and Anti-Inflammatory Effects of Zinc. Zinc-Dependent NF-ΚB Signaling. Inflammopharmacol 2017, 25, 11–24, doi:10.1007/s10787-017-0309-4.

49. Agnew, U.M.; Slesinger, T.L. Zinc Toxicity. In StatPearls; StatPearls Publishing: Treasure Island (FL), 2021.

50. Aude-Garcia, C.; Dalzon, B.; Ravanat, J.L.; Collin-Faure, V.; Diemer, H.; Strub, J.M.; Cianferani, S.; Van Dorsselaer, A.; Carriere, M.; Rabilloud, T. A Combined Proteomic and Targeted Analysis Unravels New Toxic Mechanisms for Zinc Oxide Nanoparticles in Macrophages. Journal of Proteomics 2016, 134, 174–185, doi:10.1016/j.jprot.2015.12.013.

51. Cho, W.-S.; Duffin, R.; Howie, S.E.; Scotton, C.J.; Wallace, W.A.; MacNee, W.; Bradley, M.; Megson, I.L.; Donaldson, K. Progressive Severe Lung Injury by Zinc Oxide Nanoparticles; the Role of Zn2+ Dissolution inside Lysosomes. Part Fibre Toxicol 2011, 8, 27, doi:10.1186/1743-8977-8-27.

52. Song, W.; Zhang, J.; Guo, J.; Zhang, J.; Ding, F.; Li, L.; Sun, Z. Role of the Dissolved Zinc Ion and Reactive Oxygen Species in Cytotoxicity of ZnO Nanoparticles. Toxicology Letters 2010, 199, 389–397, doi:10.1016/j.toxlet.2010.10.003.

53. Dalzon, B.; Torres, A.; Diemer, H.; Ravanel, S.; Collin-Faure, V.; Pernet-Gallay, K.; Jouneau, P.-H.; Bourguignon, J.; Cianférani, S.; Carrière, M.;, et al. How Reversible Are the Effects of Silver Nanoparticles on Macrophages? A Proteomic-Instructed View. Environmental Science: Nano 2019, 6, 3133–3157.

54. Torres, A.; Dalzon, B.; Collin-Faure, V.; Diemer, H.; Fenel, D.; Schoehn, G.; Cianférani, S.; Carrière, M.; Rabilloud, T. How Reversible Are the Effects of Fumed Silica on Macrophages? A Proteomics-Informed View. Nanomaterials (Basel) 2020, 10, 1939, doi:10.3390/nano10101939.

55. Dalzon, B.; Torres, A.; Devcic, J.; Fenel, D.; Sergent, J.-A.; Rabilloud, T. A Low-Serum Culture System for Prolonged in Vitro Toxicology Experiments on a Macrophage System. Front. Toxicol. 2021, 3, 780778, doi:10.3389/ftox.2021.780778.

56. Rabilloud, T. Optimization of the Cydex Blue Assay: A One-Step Colorimetric Protein Assay Using Cyclodextrins and Compatible with Detergents and Reducers. PLoS One 2018, 13, e0195755.

57. Abel, G.; Szollosi, J.; Fachet, J. Phagocytosis of Fluorescent Latex Microbeads by Peritoneal Macrophages in Different Strains of Mice: A Flow Cytometric Study. Eur J Immunogenet 1991, 18, 239–245.

58. Triboulet, S.; Aude-Garcia, C.; Carriere, M.; Diemer, H.; Proamer, F.; Habert, A.; Chevallet, M.; Collin-Faure, V.; Strub, J.M.; Hanau, D.;, et al. Molecular Responses of Mouse Macrophages to Copper and Copper Oxide Nanoparticles Inferred from Proteomic Analyses. Mol Cell Proteomics 2013, 12, 3108–3122.

59. Aude-Garcia, C.; Dalzon, B.; Ravanat, J.L.; Collin-Faure, V.; Diemer, H.; Strub, J.M.; Cianferani, S.; Van Dorsselaer, A.; Carriere, M.; Rabilloud, T. A Combined Proteomic and Targeted Analysis Unravels New Toxic Mechanisms for Zinc Oxide Nanoparticles in Macrophages. Journal of Proteomics 2016, 134, 174–185, doi:10.1016/j.jprot.2015.12.013.

60. Repetto, G.; del Peso, A.; Zurita, J.L. Neutral Red Uptake Assay for the Estimation of Cell Viability/Cytotoxicity. Nat Protoc 2008, 3, 1125–1131.

61. Dechézelles, J.-F.; Griffete, N.; Dietsch, H.; Scheffold, F. A General Method to Label Metal Oxide Particles with Fluorescent Dyes Using Aryldiazonium Salts. Part. Part. Syst. Charact. 2013, 30, 579–583, doi:10.1002/ppsc.201300014.

62. Triboulet, S.; Aude-Garcia, C.; Armand, L.; Collin-Faure, V.; Chevallet, M.; Diemer, H.; Gerdil, A.; Proamer, F.; Strub, J.M.; Habert, A.;, et al. Comparative Proteomic Analysis of the Molecular Responses of Mouse Macrophages to Titanium Dioxide and Copper Oxide Nanoparticles Unravels Some Toxic Mechanisms for Copper Oxide Nanoparticles in Macrophages. PLoS One 2015, 10, e0124496.

63. Armand, L.; Tarantini, A.; Beal, D.; Biola-Clier, M.; Bobyk, L.; Sorieul, S.; Pernet-Gallay, K.; Marie-Desvergne, C.; Lynch, I.; Herlin-Boime, N.;, et al. Long-Term Exposure of A549 Cells to Titanium Dioxide Nanoparticles Induces DNA Damage and Sensitizes Cells towards Genotoxic Agents. Nanotoxicology 2016, 10, 913–923, doi:10.3109/17435390.2016.1141338.

64. Kroll, A.; Pillukat, M.H.; Hahn, D.; Schnekenburger, J. Current in Vitro Methods in Nanoparticle Risk Assessment: Limitations and Challenges. Eur J Pharm Biopharm 2009, 72, 370–377, doi:10.1016/j.ejpb.2008.08.009.

65. Triboulet, S.; Aude-Garcia, C.; Armand, L.; Gerdil, A.; Diemer, H.; Proamer, F.; Collin-Faure, V.; Habert, A.; Strub, J.M.; Hanau, D.;, et al. Analysis of Cellular Responses of Macrophages to Zinc Ions and Zinc Oxide Nanoparticles: A Combined Targeted and Proteomic Approach. Nanoscale 2014, 6, 6102–6114.

66. Bourquin, J.; Septiadi, D.; Vanhecke, D.; Balog, S.; Steinmetz, L.; Spuch-Calvar, M.; Taladriz-Blanco, P.; Petri-Fink, A.; Rothen-Rutishauser, B. Reduction of Nanoparticle Load in Cells by Mitosis but Not Exocytosis. ACS Nano 2019, 13, 7759–7770, doi:10.1021/acsnano.9b01604.

67. Kluegl, I.; Hiller, K.-A.; Landthaler, M.; Baeumler, W. Incidence of Health Problems Associated with Tattooed Skin: A Nation-Wide Survey in German-Speaking Countries. Dermatology 2010, 221, 43–50, doi:10.1159/000292627.

68. Baranska, A.; Shawket, A.; Jouve, M.; Baratin, M.; Malosse, C.; Voluzan, O.; Vu Manh, T.-P.; Fiore, F.; Bajénoff, M.; Benaroch, P.;, et al. Unveiling Skin Macrophage Dynamics Explains Both Tattoo Persistence and Strenuous Removal. J Exp Med 2018, 215, 1115– 1133, doi:10.1084/jem.20171608.

69. Battistini, B.; Petrucci, F.; De Angelis, I.; Failla, C.M.; Bocca, B. Quantitative Analysis of Metals and Metal-Based Nano- and Submicron-Particles in Tattoo Inks. Chemosphere 2020, 245, 125667, doi:10.1016/j.chemosphere.2019.125667.

70. Schreiver, I.; Hutzler, C.; Laux, P.; Berlien, H.-P.; Luch, A. Formation of Highly Toxic Hydrogen Cyanide upon Ruby Laser Irradiation of the Tattoo Pigment Phthalocyanine Blue. Sci Rep 2015, 5, 12915, doi:10.1038/srep12915.

71. Bauer, E.M.; Scibetta, E.V.; Cecchetti, D.; Piccirillo, S.; Antonaroli, S.; Sennato, S.; Cerasa, M.; Tagliatesta, P.; Carbone, M. Treatments of a Phthalocyanine-Based Green Ink for Tattoo Removal Purposes: Generation of Toxic Fragments and Potentially Harmful Morphologies. Arch Toxicol 2020, 94, 2359–2375, doi:10.1007/s00204-020-02790-7.

72. Oberdörster, G. Lung Particle Overload: Implications for Occupational Exposures to Particles. Regul Toxicol Pharmacol 1995, 21, 123–135, doi:10.1006/rtph.1995.1017.

73. Warheit, D.B.; Hansen, J.F.; Yuen, I.S.; Kelly, D.P.; Snajdr, S.I.; Hartsky, M.A. Inhalation of High Concentrations of Low Toxicity Dusts in Rats Results in Impaired Pulmonary Clearance Mechanisms and Persistent Inflammation. Toxicol Appl Pharmacol 1997, 145, 10– 22, doi:10.1006/taap.1997.8102.

74. Wiemann, M.; Vennemann, A.; Sauer, U.G.; Wiench, K.; Ma-Hock, L.; Landsiedel, R. An in Vitro Alveolar Macrophage Assay for Predicting the Short-Term Inhalation Toxicity of Nanomaterials. J Nanobiotechnol 2016, 14, 16, doi:10.1186/s12951-016-0164-2.

75. Tsuboi, A.; Kurotsu, T.; Terasima, T. Changes in Protein Content per Cell during Growth of Mouse L Cells. Exp Cell Res 1976, 103, 257–261, doi:10.1016/0014-4827(76)90262-7.

76. Carriere, M.; Arnal, M.-E.; Douki, T. TiO2 Genotoxicity: An Update of the Results Published over the Last Six Years. Mutat Res 2020, 854–855, 503198, doi:10.1016/j.mrgentox.2020.503198.

77. Carriere, M.; Sauvaigo, S.; Douki, T.; Ravanat, J.-L. Impact of Nanoparticles on DNA Repair Processes: Current Knowledge and Working Hypotheses. MUTAGE 2017, 32, 203– 213, doi:10.1093/mutage/gew052.

78. Dalzon, B.; Aude-Garcia, C.; Diemer, H.; Bons, J.; Marie-Desvergne, C.; Pérard, J.; Dubosson, M.; Collin-Faure, V.; Carapito, C.; Cianférani, S.;, et al. The Longer the Worse: A Combined Proteomic and Targeted Study of the Long-Term versus Short-Term Effects of Silver Nanoparticles on Macrophages. Environ. Sci.: Nano 2020, 7, 2032–2046, doi:10.1039/C9EN01329F.

79. Deshmane, S.L.; Kremlev, S.; Amini, S.; Sawaya, B.E. Monocyte Chemoattractant Protein-1 (MCP-1): An Overview. Journal of Interferon & Cytokine Research 2009, 29, 313– 326, doi:10.1089/jir.2008.0027.

80. Bhavsar, I.; Miller, C.S.; Al-Sabbagh, M. Macrophage Inflammatory Protein-1 Alpha (MIP-1 Alpha)/CCL3: As a Biomarker. In General Methods in Biomarker Research and their Applications; Preedy, V.R., Patel, V.B., Eds.; Biomarkers in Disease: Methods, Discoveries and Applications; Springer Netherlands: Dordrecht, 2015; pp. 223–249 ISBN 978-94-007-7695-1.

81. Vaddi, K.; Newton, R.C. Regulation of Monocyte Integrin Expression by Beta-Family Chemokines. J Immunol 1994, 153, 4721–4732.

82. Gschwandtner, M.; Derler, R.; Midwood, K.S. More Than Just Attractive: How CCL2 Influences Myeloid Cell Behavior Beyond Chemotaxis. Front. Immunol. 2019, 10, 2759, doi:10.3389/fimmu.2019.02759.

83. Grove, G.L.; Kligman, A.M. Age-Associated Changes in Human Epidermal Cell Renewal. Journal of Gerontology 1983, 38, 137–142, doi:10.1093/geronj/38.2.137.

84. Vandendriessche, S.; Cambier, S.; Proost, P.; Marques, P.E. Complement Receptors and Their Role in Leukocyte Recruitment and Phagocytosis. Front. Cell Dev. Biol. 2021, 9, 624025, doi:10.3389/fcell.2021.624025.

85. Ehirchiou, D.; Xiong, Y.; Xu, G.; Chen, W.; Shi, Y.; Zhang, L. CD11b Facilitates the Development of Peripheral Tolerance by Suppressing Th17 Differentiation. Journal of Experimental Medicine 2007, 204, 1519–1524, doi:10.1084/jem.20062292.

86. Han, C.; Jin, J.; Xu, S.; Liu, H.; Li, N.; Cao, X. Integrin CD11b Negatively Regulates TLR-Triggered Inflammatory Responses by Activating Syk and Promoting Degradation of MyD88 and TRIF via Cbl-b. Nat Immunol 2010, 11, 734–742, doi:10.1038/ni.1908.

87. Varga, G.; Balkow, S.; Wild, M.K.; Stadtbaeumer, A.; Krummen, M.; Rothoeft, T.; Higuchi, T.; Beissert, S.; Wethmar, K.; Scharffetter-Kochanek, K.;, et al. Active MAC-1 (CD11b/CD18) on DCs Inhibits Full T-Cell Activation. Blood 2007, 109, 661–669, doi:10.1182/blood-2005-12-023044.

88. Zhang, Q.; Lee, W.-B.; Kang, J.-S.; Kim, L.K.; Kim, Y.-J. Integrin CD11b Negatively Regulates Mincle-Induced Signaling via the Lyn-SIRPα-SHP1 Complex. Exp Mol Med 2018, 50, e439, doi:10.1038/emm.2017.256.

89. Schmid, M.C.; Khan, S.Q.; Kaneda, M.M.; Pathria, P.; Shepard, R.; Louis, T.L.; Anand, S.; Woo, G.; Leem, C.; Faridi, M.H.;, et al. Integrin CD11b Activation Drives Anti-Tumor Innate Immunity. Nat Commun 2018, 9, 5379, doi:10.1038/s41467-018-07387-4.

90. Díez-Tercero, L.; Delgado, L.M.; Bosch-Rué, E.; Perez, R.A. Evaluation of the Immunomodulatory Effects of Cobalt, Copper and Magnesium Ions in a pro Inflammatory Environment. Sci Rep 2021, 11, 11707, doi:10.1038/s41598-021-91070-0.

