## Supplementary material for "Immediate and sustained effects of cobalt and zinc-containing pigments on macrophages": supdata for manuscript

### Supplementary data:

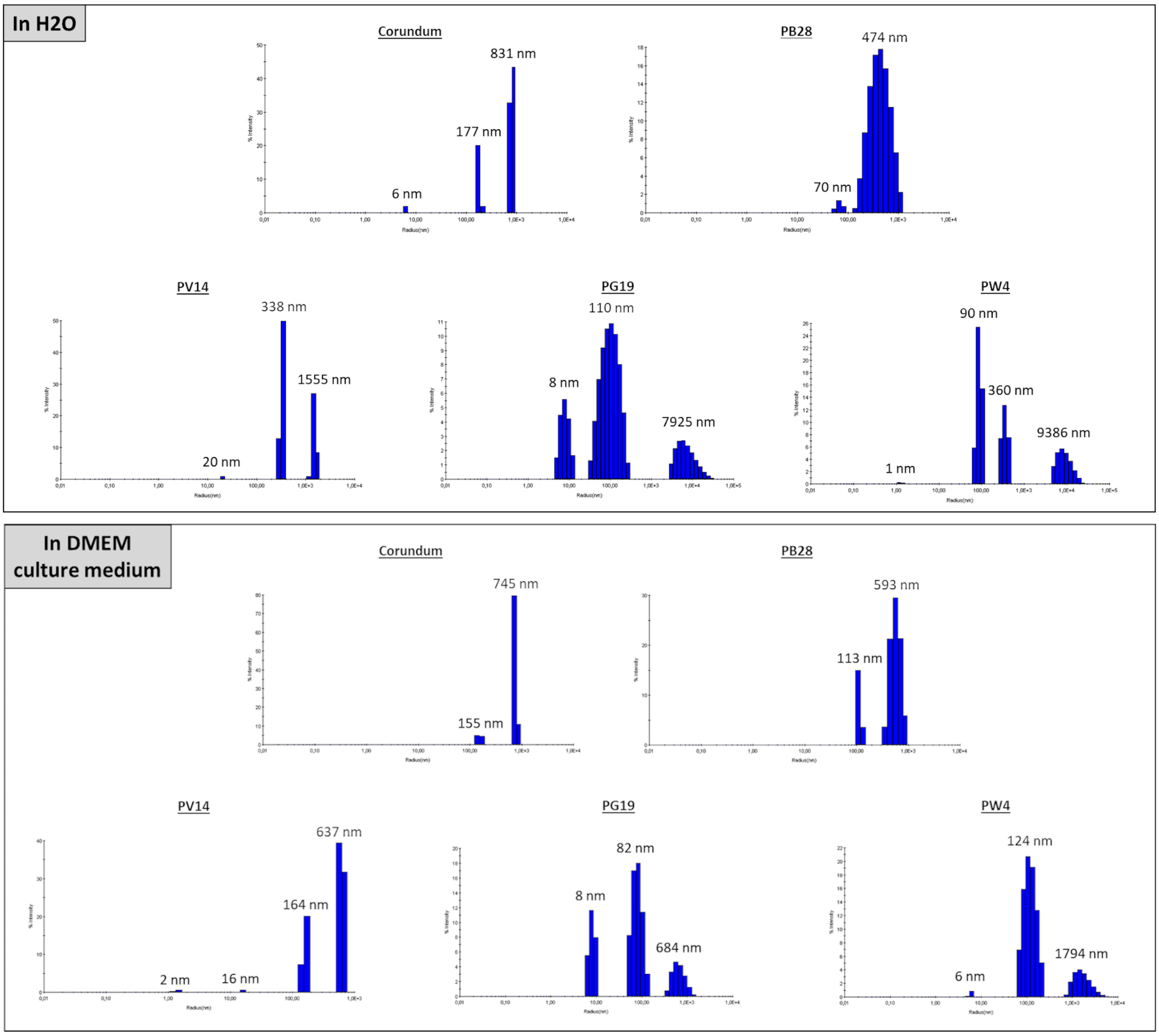


*Supplementary data 1 – Example of hydrodynamic diameter measurement (screen print of raw data) by DLS in $H_2O$ and DMEM culture medium.*

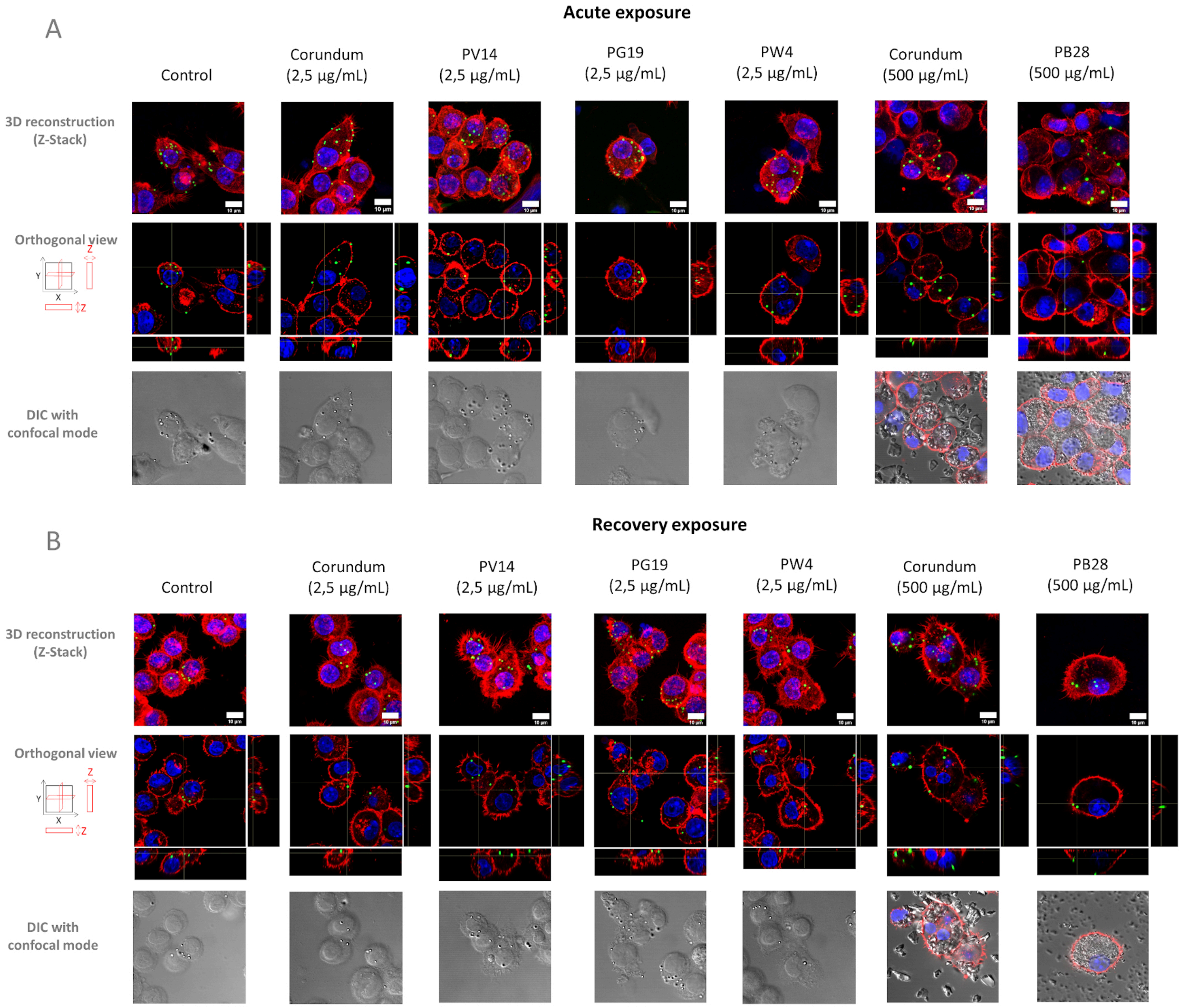


*Supplementary data 2 – Visualization of phagocytic activity by confocal microscopy. Pannel A- Acute exposure of macrophages to pigments. Pannel B – End of recovery exposure of macrophages to pigment. Red color = Actin filament visualized with phalloidin-Atto 560. Blue color = Nucleus visualized with Dapi. Green color = Fluorescent yellow/green carboxylate-modified-polystyren beads (1 µm diameter). Grey color = differential interference contrast (DIC). Scale bar = 10 µm.*

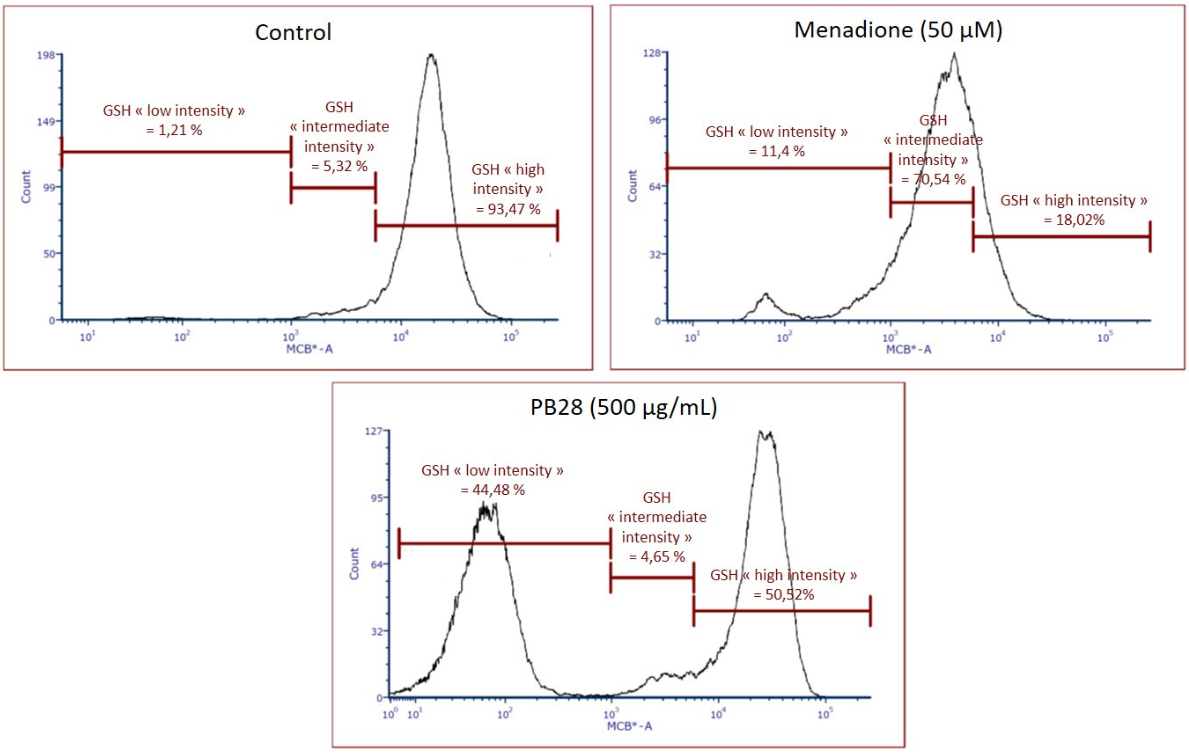


*Supplementary data 3 – Examples of GSH results obtained via flow cytometry (Raw results of recovery exposure).*

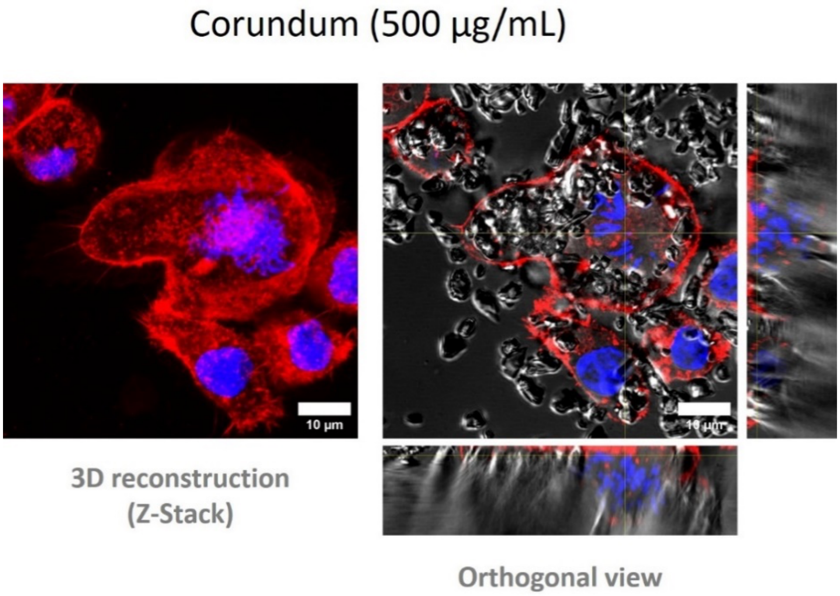


*Supplementary data 4 – Confocal microscopy. Observation of actin conformation and nuclear integrity. Red = Actin (in red) stained with phalloidin-Atto 560. DNA (in blue) stained with Dapi. Grey = Corundum particles observed in confocal DIC. Scale bar = 10 µm.*
